# Humanized V(D)J-rearranging and TdT-expressing Mouse Vaccine Models with Physiological HIV-1 Broadly Neutralizing Antibody Precursors

**DOI:** 10.1101/2022.10.12.511911

**Authors:** Sai Luo, Changbin Jing, Adam Yongxin Ye, Sven Kratochvil, Christopher A. Cottrell, Ja-Hyun Koo, Aimee Chapdelaine Williams, Lucas Vieira Francisco, Himanshu Batra, Edward Lamperti, Oleksandr Kalyuzhniy, Yuxiang Zhang, Alessandro Barbieri, John P. Manis, Barton F. Haynes, William R. Schief, Facundo D. Batista, Ming Tian, Frederick W. Alt

**Author notes:** These authors contributed equally to this work (Sai Luo, Changbin Jing, Adam Yongxin Ye). Correspondence: Frederick W. Alt or Ming Tian or. **Author Contributions:** S.L., M.T., and F.W.A. designed the experiments. S.L. generated the mouse model. M.T. and S.L. made the immunogens. S.L. and E.L. performed immunizations. C.J., S.L., S.K. and J.K. characterized antibodies. C.A.C. and O.K. expressed the antibodies and measured the kinetics and affinity of antibodies to eOD-GT8. A.B., J.P.M. and Y.Z. isolated the human tonsil naïve B cells and extracted genomic DNA. A.Y.Y. performed the bioinformatics analyses for CDR3 diversity and microhomology-mediated end joining for all experiments shown. A.C.W. performed ES cell injections. S.L., L.V.F., E.L. and H.B. performed mouse maintenance. S.L., M.T., and F.W.A. designed figures and drafted the manuscript. B.F.H., W.R.S., C.J., A.Y.Y. and F.D.B contributed to polishing the manuscript.

## Abstract

Antibody heavy chain (HC) and light chain (LC) variable region exons are assembled by V(D)J recombination. V(D)J junctional regions encode complementarity-determining-region 3 (CDR3), an antigen-contact region immensely diversified through non-templated nucleotide additions (“N-regions”) by terminal deoxynucleotidyl transferase (TdT). HIV-1 vaccine strategies seek to elicit human HIV-1 broadly neutralizing antibodies (bnAbs), such as the potent CD4-binding site VRC01-class bnAbs. Mice with primary B cells that express receptors (BCRs) representing bnAb precursors are used as vaccination models. VRC01-class bnAbs uniformly use human HC V_H_1-2 and commonly use human LCs Vκ3-20 or Vκ1-33 associated with an exceptionally short 5-amino-acid (5-aa) CDR3. Prior VRC01-class models had non-physiological precursor levels and/or limited precursor diversity. Here, we describe VRC01-class rearranging mice that generate more physiological primary VRC01-class BCR repertoires via rearrangement of V_H_1-2, as well as Vκ1-33 and/or Vκ3-20 in association with diverse CDR3s. Human-like TdT expression in mouse precursor B cells increased LC CDR3 length and diversity and also promoted generation of shorter LC CDR3s via N-region suppression of dominant microhomology-mediated Vκ-to-Jκ joins. Priming immunization with eOD-GT8 60mer, which strongly engages VRC01 precursors, induced robust VRC01-class germinal center (GC) B cell responses. Vκ3-20-based responses were enhanced by N-region addition, which generates Vκ3-20-to-Jκ junctional sequence combinations that encode VRC01-class 5-aa CDR3s with a critical E residue. VRC01-class-rearranging models should facilitate further evaluation of VRC01-class prime and boost immunogens. These new VRC01-class mouse models establish a prototype for generation of vaccine-testing mouse models for other HIV-1 bnAb lineages that employ different HC or LC Vs.

**Significance Statement:** Mouse models that express human precursors of HIV-1 broadly neutralizing antibodies (bnAbs) are useful for evaluating vaccination strategies for eliciting such bnAbs in humans. Prior models were handicapped by non-physiological frequency and/or diversity of B lymphocytes that express the bnAb precursors. We describe a new class of mouse models in which the mice express humanized bnAb precursors at a more physiologically relevant level through developmental rearrangement of both antibody heavy and light chain gene segments that encode the precursors. The model also incorporated a human enzyme that diversifies the rearranging gene segments and promotes generation of certain variable region sequences needed for the response. This new class of mouse models should facilitate preclinical evaluation of candidate human HIV-1 vaccination strategies.

## Introduction

Diverse antibody variable region exons are assembled in developing B cells from Immunoglobulin (Ig) HC V, D, and J gene segments and from Igκ or Igλ LC V and J segments (1). In humans, there are 55 germline HC Vs (V_H_s) and 70 Igκ and Igλ LC Vs. Vs encode most of the HC and LC variable region, including the antigen contact CDR1 and CDR2 sequences that vary among different HC and LC Vs. Ig HC V(D)J recombination occurs at the progenitor (Pro) B cell developmental stage in the fetal liver and in the postnatal bone marrow (2, 3). Ig LC V to J recombination takes place in the subsequent precursor (Pre) B cell developmental stage in these same sites (1). T cell receptor variable region exon assembly also occurs in the fetal liver and thymus and then in the postnatal thymus (4, 5). Mice also have similar sets of Ig HC and LC and TCR variable region gene segments as those found in humans and, in general, assemble them in the context of similar developmental processes (6, 7).

Primary BCR diversity is achieved, in part, by assorting HC and LC Vs along with each of their distinct sets of CDR1 and 2 sequences. However, several V(D)J junctional diversification mechanisms play an even greater role in V(D)J diversity generation (8). In this regard, TdT, a DNA polymerase that adds nucleotides to 3’DNA ends without a template (9), plays a key role. In this regard, V(D)J junctional diversity is immensely augmented by TdT-based non-templated nucleotide additions, referred to as N regions (10), that are added to V(D)J junctions. While N-region addition generates CDR3 length and sequence diversity, it also suppresses recurrent CDR3s resulting from microhomology (MH)-mediated V(D)J joining (10–13). TdT expression is absent during fetal B and T cell development, resulting in less diverse repertoires dominated by variable region exons promoted by recurrent MH-mediated joins (14–21). In contrast, TDT expression diversifies antigen receptor variable region repertoires generated in mouse and human developing B and T cells that develop postnatally, with the notable exception of LC variable region repertoires in mice (10, 22, 23). Thus, while TdT is expressed during LC V(D)J recombination in postnatal human Pre-B cells (24), it is not expressed in postnatal mouse pre-B cells (25, 26), leading to decreased junctional diversity and much more abundant MH-mediated joins in primary mouse LC repertoires compared to those of humans (22, 23). Lack of TdT expression in fetal repertoires also is known to promote recurrent MH-mediated V(D)J junctions, that are not dominant in post-natal repertoires due to TdT expression. Some such recurrent MH-mediated V(D)J joins in fetal T or B cell repertoires generate TCRs or BCRs critical for certain physiological responses (13, 14, 27, 28). However, the potential role of TdT and N regions in promoting specific responses has remained largely unaddressed.

VRC01-class bnAb HCs employ human V_H_1-2, which encodes residues that contact the HIV-1 envelope protein (Env) CD4 binding site (29–37). VRC01-class LC variable regions are known to be encoded by several Vs; but all are associated with an exceptionally short 5 amino acid (5-aa) CDR3, which avoids steric clash with Env and contributes to Env interaction (29–37). As both requirements can be achieved by V(D)J recombination, they are predicted attributes of primary VRC01-class precursor BCRs. However, inferred primary VRC01-class BCRs lack detectable affinity for naïve Envs (38–41). In this regard, following BCR antigen-activation, primary B cells are driven into GC reactions where they undergo rounds of variable region exon somatic hyper-mutation (SHM) followed by selection of SHMs that increase BCR antigen-binding affinity. This process ultimately leads to high-affinity antibody production. Correspondingly, a third VRC01-class bnAb attribute is abundant variable region SHMs with only a subset contributing to broad Env-binding and potent VRC01-class bnAb activity (37, 42), consistent with VRC01-class bnAb evolution occurring over long HIV-1 infection times and many SHM/selection cycles.

To elicit VRC01-class bnAbs, sequential vaccine immunization approaches propose a priming immunogen to drive precursors into GCs followed by boost immunogens designed to lead them through rounds of SHM/affinity maturation. Based on a structurally designed eOD-GT8 immunogen that binds to the inferred VRC01 unmutated common ancestor (UCA) BCR, potential human VRC01-like precursor B cell frequency was estimated to be 1 in 400,000 or fewer (43, 44). To test priming and sequential immunogens that could elicit VRC01-class bnAbs in humans, mouse models are needed that reflect as closely as possible the biology of human B cell responses. Early models expressed knock-in V_H_1-2 HCs and, in some, VRC01-class LC Vs, both with mature CDR3s (45–47). These models were non-physiologic as their BCR repertoire was dominated by a single human HC/LC combination or a single human HC with diverse mouse LCs. Mice with fully human HC and LC gene segment loci assembled by V(D)J recombination were also tested; but precursor frequencies were 150- to 900-fold lower than that of humans (48), likely due to inability to express immense human-like CDR3 repertoires in mice with orders of magnitude fewer B cells. A V_H_1-2-rearranging mouse model generated diverse V_H_1-2 HC CDR3s, but it employed a germline-reverted VRC01 precursor LC with a 5-aa CDR3 from mature VRC01 bnAb (49). While useful for HC maturation studies during sequential immunization, this model was limited by over-abundance of VRC01 lineage LC precursors. More recently, B cells from transgenic VRC01-class UCA or eOD-GT8-binding precursor knock-in mice were adoptively transferred into congenic recipient mice at human-like frequencies (50–53). While this elegant approach has been very useful, it still has certain limitations as it focused only on eOD-GT8-priming and tested just a small subset of potential VRC01 lineage precursors (50–53).

## Results

### Generation of mice with VRC01-class-rearranging human HC and LC Vs

To address issues of prior models, we developed complete VRC01 mouse models in which individual B cells express one of a multitude of different VRC01 precursors at human-like frequencies, based on enforced rearrangement of both V_H_1-2 and VRC01-class Vκs (Fig.1A). All complete VRC01-class models employ our previously described V_H_1-2-rearranging HC allele in which the most D proximal functional mouse V_H_ (V_H_81X) was replaced with human V_H_1-2 (49, 54). The CTCF-binding site (CBE)-based IGCR1 element in the V_H_ to D interval is also inactivated on this allele, which leads to dominant rearrangement of human V_H_1-2 in an otherwise intact upstream mouse V_H_ locus (55). On this allele, high-level V_H_1-2 utilization in the absence of IGCR1 is mediated by its closely associated downstream CBE element (56). Our new models also use a version of this rearranging HC allele in which the mouse J_H_ segments were replaced with human J_H_2, which can contribute a trptophan residue (Trp100B) conserved in the HC CDR3 of VRC01-class bnAbs (54). We have retained mouse Ds in the model for reasons we have previously described (57). On homozygous replacement alleles in our new VRC01-class models, V_H_1-2 rearrangements represent nearly 73.8% of primary V(D)J rearrangements (Fig. S1A, upper panel). Due to counter selection of lower frequency upstream mouse V_H_ rearrangements, V_H_1-2 contribution to primary B cell BCR repertoires is reduced to 43% (Fig. S1A, bottom panel), with immense CDR3 diversity (Fig. S1B). Such CDR3 diversity is critical, as V_H_1-2-encoded HC CDR3s were implicated in Env recognition by precursor VRC01-class BCRs and also implicated in maturation of VRC01-class bnAbs (58, 59).

**Figure 1.**
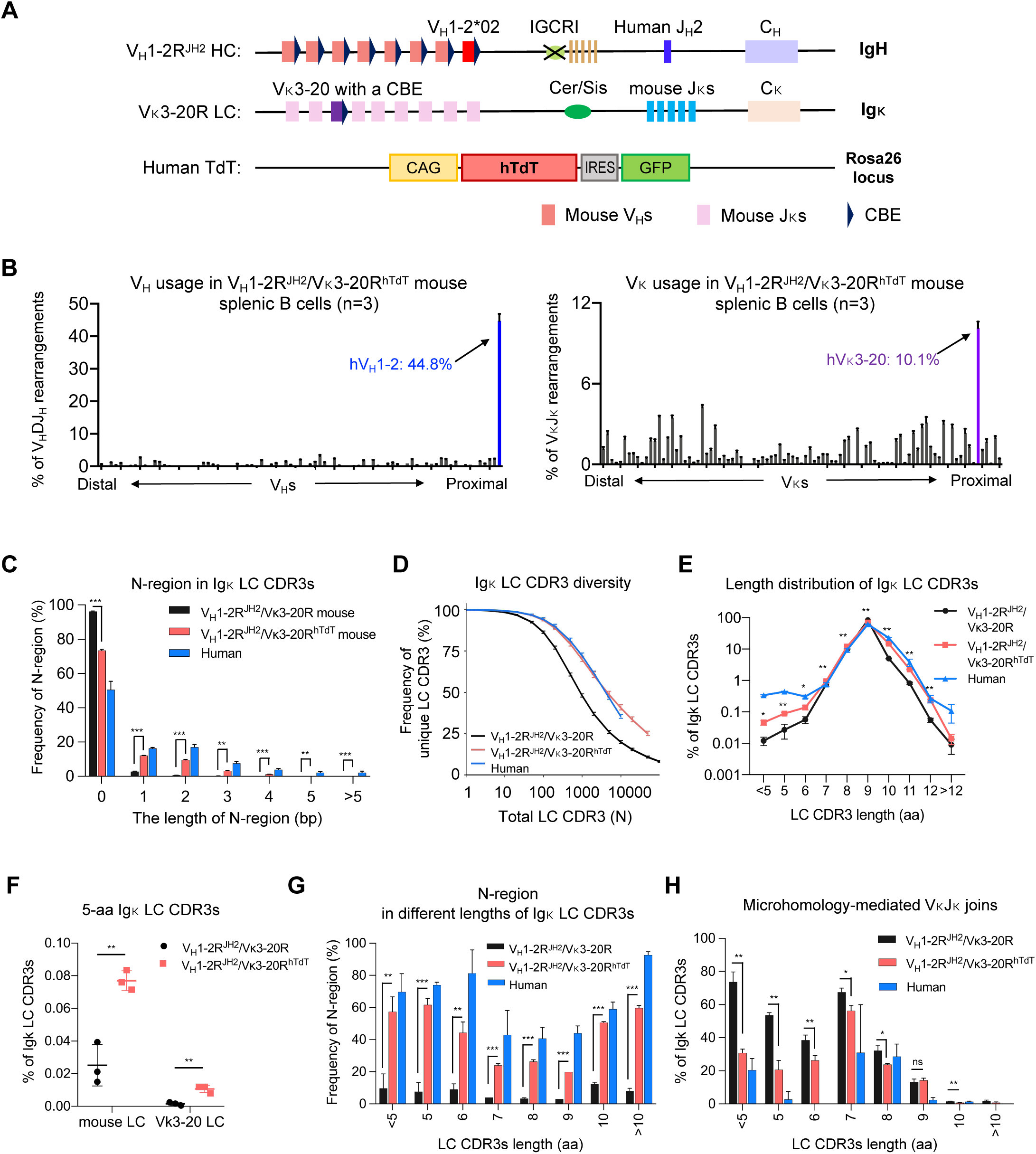
Generation and characterization of the V_H_1-2^JH2^/Vκ3-20^hTdT^-rearranging mouse models. (A) Illustration of genetic modifications in the *Igh* and *Igκ*locus of V_H_1-2^JH2^/Vκ3-20-rearranging mouse models. The most D_H_-proximal functional mouse V_H_ (V_H_81X) was replaced with the human V_H_1-2 on an IGCR1-deleted allele. The mouse J_H_s were replaced with the human J_H_2. The *Jκ*-proximal Vκ3-7 was replaced with human Vκ3-20 plus a CBE 50bp downstream of its RSS. Human TdT gene was knocked into mouse *Rosa* locus. (B) HTGTS-rep-seq analysis of V_H_ (upper panel) and Vκ (bottom panel) usage in V_H_1-2^JH2^/Vκ3-20^hTdT^-rearranging mouse splenic B cells. The x axis listed all functional V_H_s or Vκs from the distal to the *D-* or *Jκ*-proximal end. The histogram displayed the percent usage of each V_H_ or Vκs among all productive V_H_DJ_H_ or VκJκ rearrangements. The usage of human V_H_1-2 and Vκ3-20 were shown in blue and purple, respectively. (C) Length distribution of N regions in VκJκ junctions from human, V_H_1-2R^JH2^/Vκ3-20R mouse and V_H_1-2R^JH2^/Vκ3-20R^hTdT^ mouse naïve B cells. The human naïve B cells were isolated from human tonsils using CD19^+^, IgD^+^, CD27^-^ and CD38^-^. (D) The diversity of Igκ LC CDR3s in human, VH1-2R^JH2^/Vκ3-20R mouse and VH1-2R^JH2^/Vκ3-20R^hTdT^ mouse naive B cells. The x axis represents the total Igκ LC CDR3 number (N). The y axis represents the frequency of unique Igκ LC CDR3s among total Igκ LC CDR3s. The differences of CDR3 diversities between VH1-2R^JH2^/Vκ3-20R and VH1-2R^JH2^/Vκ3-20R^hTdT^ mice are significant when the total CDR3 number is above 50 (*p*<0.001 for N>=50). (E) Length distribution of Igκ LC CDR3s in human, VH1-2R^JH2^/Vκ3-20R mouse and VH1-2R^JH2^/Vκ3-20R^hTdT^mouse naïve B cells. (F) The frequency of 5-aa LC CDR3 in VH1-2R^JH2^/Vκ3-20R and VH1-2R^JH2^/Vκ3-20R^hTdT^ mouse naïve B cells. (G) Frequency of N regions in different length of Igκ LC CDR3s from human, VH1-2R^JH2^/Vκ3-20R and VH1-2R^JH2^/Vκ3-20R^hTdT^ mouse naïve B cells. (H) Frequency of MH-mediated VκJκ joins in Igκ LC CDR3s from human, VH1-2R^JH2^/Vκ3-20 and VH1-2R^JH2^/Vκ3-20R^hTdT^ mouse naïve B cells. Data from (B), (C), (E), (F), (G) and (H) were mean ± SD of three independent experiments. Statistical comparisons in (C), (E), (F), (G) and (H) were performed between VH1-2R^JH2^/Vκ3-20R and VH1-2R^JH2^/Vκ3-20R^hTdT^ mice using a two-tailed unpaired t test. \**p* <0.05, \*\**p* <0.01, \*\*\**p* <0.001

To generate human Vκ-rearranging LC alleles, we used a strategy similar to that which we used for V_H_1-2, as recently described (57). The CBE-based Cer/Sis element in the Vκ to Jκ interval has been implicated in promoting distal versus proximal Vκ rearrangements (60). To test Cer/Sis functions in more detail, we deleted this element from the wild-type mouse allele and assessed impact on Vκ rearrangement via our high throughput HTGTS-Rep-seq method (Fig. S2A). Homozygous Cer/Sis deletion substantially increased (up to 8-fold) the frequency of 7 of the 11 the most Jκ-proximal Vκs (Fig.S2B). Indeed, these 7 Vκs contributed to the vast majority of the primary BCR repertoire of these mice (Fig.S2C), as upstream Vκ rearrangements were essentially abrogated in the absence of Cer/Sis. We note that Vκ3-2 and Vκ3-7 showed the greatest increase in utilization in the absence of Cer/Sis. Our initial plan for our VRC01-rearranging mouse models, analogous to our V_H_1-2 rearranging *Igh* allele (49), was to increase utilization of human Vκs in the model by introducing them into proximal positions on Cer/Sis-deleted *Igκ* alleles.

We replaced the Vκ3-2 sequence encoding the leader-intron-V sequence with the corresponding sequences of human Vκ1-33 on a wild-type *Igκ* allele (“Vκ1-33-rearranging” allele) and then also deleted Cer/Sis on that allele (“Vκ1-33^CSΔ^-rearranging” allele) (57). In these replacement alleles, we maintained the mouse Vκ3-2 sequence upstream of the ATG (including the promoter) and the Vκ3-2 downstream sequence starting at the Vκ3-2 RSS. HTGTS-Rep-seq revealed that, similarly to Vκ3-2, human Vκ1-33 on homozygous replacement alleles in our VRC01-class models accounted for approximately 2% or 17% of primary Vκ rearrangements in presence or absence of Cer/Sis element, respectively (Fig. S3A). Vκ1-33 contributed to the splenic BCR repertoire at similar frequencies (approximately 2% and 15%, respectively; Fig. S3B). We also generated a “Vκ3-20-rearranging allele” in which mouse proximal Vκ3-7 was replaced with human Vκ3-20 (Fig. 1A; Fig. S3, C and D). When homozygous in mice, the Vκ3-20-rearranging allele contributed about 6% of primary Vκ rearrangements and contributed similar frequencies in splenic BCR repertoires (Fig. S3E). We considered these levels sufficiently high to leave Cer/Sis intact for initial experiments.

Based on studies of the *Igh* locus (56), we also inserted CBEs just downstream of the RSSs of the inserted Vκ1-33 and Vκ3-20 gene segments (Fig. 1A) (57). However, we found that, compared to the rearrangement frequencies of mouse Vκs they replaced, inserted CBEs had no measurable effect on Vκ1-33 rearrangement either in the presence or absence of Cer/Sis (Fig.S2C; Fig.S3, A and B) and only modestly increased Vκ3-20 rearrangement in the presence of Cer/Sis (Fig.S2C; Fig.S3E). These findings, particularly, the lack of the attached CBE to dominantly increase Vκ1-33 rearrangement in the absence of Cer/Sis, suggest that mechanisms underlying CBE-enhanced dominant utilization of proximal V_H_s in the absence of IGCR1 may not conserved in the context of Igκ V(D)J recombination. This notion is consistent with recent findings, published after these models were generated, that indicated mechanisms that promote long-range V_H_ to DJ_H_ joining are, at least in part, distinct from those that promote long-range Vκ to Jκ joining (61).

We refer to these new VRC01-class mouse models with human V_H_1-2- and Vκ-rearranging (“R”) alleles as the V_H_1-2R^JH2^/Vκ1-33R model, the V_H_1-2R^JH2^/Vκ1-33R^CSΔ^ model (^”CSΔ”^ indicates Cer/Sis deletion), and the V_H_1-2R^JH2^/Vκ3-20R model. Based on fluorescence-activated cell sorting (FACS) analyses of cell surface markers, splenic B and T cell populations in all three models were comparable to those of wild-type mice (Fig. S3F). During our studies of the V_H_1-2R^JH2^/Vκ3-20R model, we discovered that the inserted Vκ3-20 sequence had acquired a single in-frame point mutation in CDR1 that changes an S to I residue (AGT to ATT) (Fig. S4A). We then corrected this mutation in the Vκ3-20 allele, introduced it into all mouse models described, and repeated all experiments originally performed with the mutated allele with mouse models harboring the corrected allele. Based on fluorescence-activated cell sorting (FACS) analyses of cell surface markers, splenic B and T cell populations in the Vκ3-20 corrected model were also comparable to those of wild-type mice and those of the mouse models harboring mutated Vκ3-20 sequence (Fig. S3F). Indeed, in all experiments described below, mouse models harboring the mutated and corrected Vκ3-20 sequence gave very similar results with respect to Vκ3-20-based VRC01-class responses, which, for comparison, are included in all immunization experiments and related figures described below.

### Enforced human TdT Expression diversifies LC repertoires

VRC01-class bnAb LCs commonly have a LC 5-aa CDR3 with a relatively conserved QQYEF amino acid sequence (32, 62). However, as compared to the frequency of LC 5-aa CDR3s in human BCR repertoires, our initial VRC01-class mouse models had 20- to 50-fold lower frequencies of LC 5-aa CDR3s (0.02%) in their mouse Vκ and human Vκ1-33 or Vκ3-20 LC BCR repertoires (Fig. S5A) (48, 62). In this regard, approximately 80% of human LC 5-aa-CDR3s are encoded by sequences with hTdT-generated N regions (Fig. S5B). Thus, to enforce more human-like TdT expression in mouse bone marrow precursor B cells which normally lack TdT expression, we targeted human hTdT into the *Rosa* locus of ES cells containing the Vκ3-20R allele (Fig. 1A; Fig.S5, C and D); as *Rosa* and *Igκ* both lie on chromosome 6, these two modifications are linked in subsequent crosses. Mice harboring the resulting Vκ3-20R^hTdT^ modified chromosome were bred to homozygosity with the V_H_1-2R^JH2^ allele to create V_H_1-2R^JH2^/Vκ3-20R^hTdT^ mice. The V_H_1-2R^JH2^/Vκ3-20R^hTdT^ mice indeed now expressed human TdT in their progenitor and precursor B cell population (Fig.S5, E and F). HTGTS-Rep-seq revealed that enforced TdT expression modestly increased Vκ3-20 expression and had little impact on utilization of V_H_1-2 in splenic B cell populations (Fig. 1B and Fig. S5G).

As compared to splenic B cells of V_H_1-2R^JH2^/Vκ3-20R mice, those of V_H_1-2R^JH2^/Vκ3-20R^hTdT^ mice had markedly increased frequencies of N regions in both mouse Vκ to Jκ junctions and human Vκ3-20 to Jκ junctions (Fig. 1C), and, correspondingly, much more diverse CDR3s (Fig. 1D). Notably, while enforced N region addition increased the proportion of longer LC CDR3s (> 9-aa), it also increased, up to 5-fold, the proportion of short mouse and Vκ3-20 LC CDR3s (< 7-aa), including 5-aa CDR3s (Fig. 1, E and F). Correspondingly, the proportion of N-regions in short LC CDR3s was significantly increased (Fig. 1G) and the proportion of MH-mediated short Vκ to Jκ joins (< 7-aa) was significantly reduced in splenic B cells of V_H_1-2R^JH2^/Vκ3-20R^hTdT^ mice as compared to those of V_H_1-2R^JH2^/Vκ3-20R mice (Fig. 1H). In addition, we compared the LC CDR3s in splenic B cells of V_H_1-2R^JH2^/Vκ3-20R and V_H_1-2R^JH2^/Vκ3-20R^hTdT^ mice to those in human tonsil naïve B cells and found that enforced TdT expression in V_H_1-2R^JH2^/Vκ3-20R^hTdT^ mice yielded more human-like CDR3s (Fig. 1, C to E, G and H). As endogenous mouse TdT expression is already robust in V_H_1-2R^JH2^/Vκ3-20R progenitor-stage B cells that undergo HC locus V(D)J recombination, human TdT expression had no obvious effect on HC CDR3 length and diversity in V_H_1-2R^JH2^/Vκ3-20R^hTdT^ mice (Fig. S5H).

We similarly introduced hTdT into the *Rosa* locus of V_H_1-2R^JH2^/Vκ1-33R^CSΔ^ mice and generated V_H_1-2R^JH2^/Vκ1-33R^CSΔ/hTdT^ mice. Analyses of splenic B cells from these two models revealed little effect of enforced hTdT expression on overall Vκ1-33 utilization and V_H_1-2 utilization in splenic B cell populations (Fig.S6A). However, as in the V_H_1-2R^JH2^/Vκ3-20R^hTdT^ model, Vκ1-33 LC CDR3 diversity and the frequency of Vκ1-33 5-aa CDR3s were significantly increased after hTdT expression (Fig.S6, B and C).

### Human TdT enhanced VRC01-class GC responses induced by eOD-GT8

To test if the human TdT expression affects the VRC01-class GC response, we immunized V_H_1-2R^JH2^/Vκ3-20R, V_H_1-2R^JH2^/Vκ1-33R^CSΔ^, V_H_1-2R^JH2^/Vκ3-20R^hTdT^ and V_H_1-2R^JH2^/Vκ1-33R^CSΔ/hTdT^ mice with eOD-GT8 60mer and poly I:C adjuvant (Fig.2A). All mice developed CD4-binding site (CD4bs)-specific germinal center (GC) responses by day 8 post-immunization, as demonstrated by the presence of GC B cells that bound eOD-GT8 but not ΔeOD-GT8 (which is a VRC01-class epitope knockout variant) (Fig. S7, A to C). We flow-sorted eOD-GT-specific GC B cells and sequenced their BCRs (Fig. 2B). We refer to B cells with VRC01-class BCRs (V_H_1-2 HCs and LCs with 5-aa CDR3s) as VRC01/Vκ1-33, VRC01/Vκ3-20 and VRC01/mVκ B cells, according to the LC they express. At day 8, VRC01/Vκ3-20 and VRC01/mVκ represented 5% and 4%, respectively of CD4bs-specific GC B cells in V_H_1-2R^JH2^/Vκ3-20R mice and 28% and 20%, respectively in V_H_1-2R^JH2^/Vκ3-20R^hTdT^ mice (Fig.2, B and C). Thus, enforced TdT expression in V_H_1-2R^JH2^/Vκ3-20R line increases frequency of VRC01-class GC B cells by approximately 5-fold. At day 8, VRC01/Vκ1-33 GC B cells represented up to 70% of CD4bs-specific GC B cells in both V_H_1-2R^JH2^/Vκ1-33R^CSΔ^ and V_H_1-2R^JH2^/Vκ1-33R^CSΔ/hTdT^ mice but no mouse VRC01/mVκs B cells were observed (Fig.2, B and C). The lack of mouse VRC01/mVκs B cells in the GCs of immunized V_H_1-2R^JH2^/Vκ1-33R^CSΔ^, V_H_1-2R^JH2^/Vκ1-33R^CSΔ/hTdT^ mice probably results from domination of the response by VRC01/Vκ1-33 B cells.

**Figure 2.**
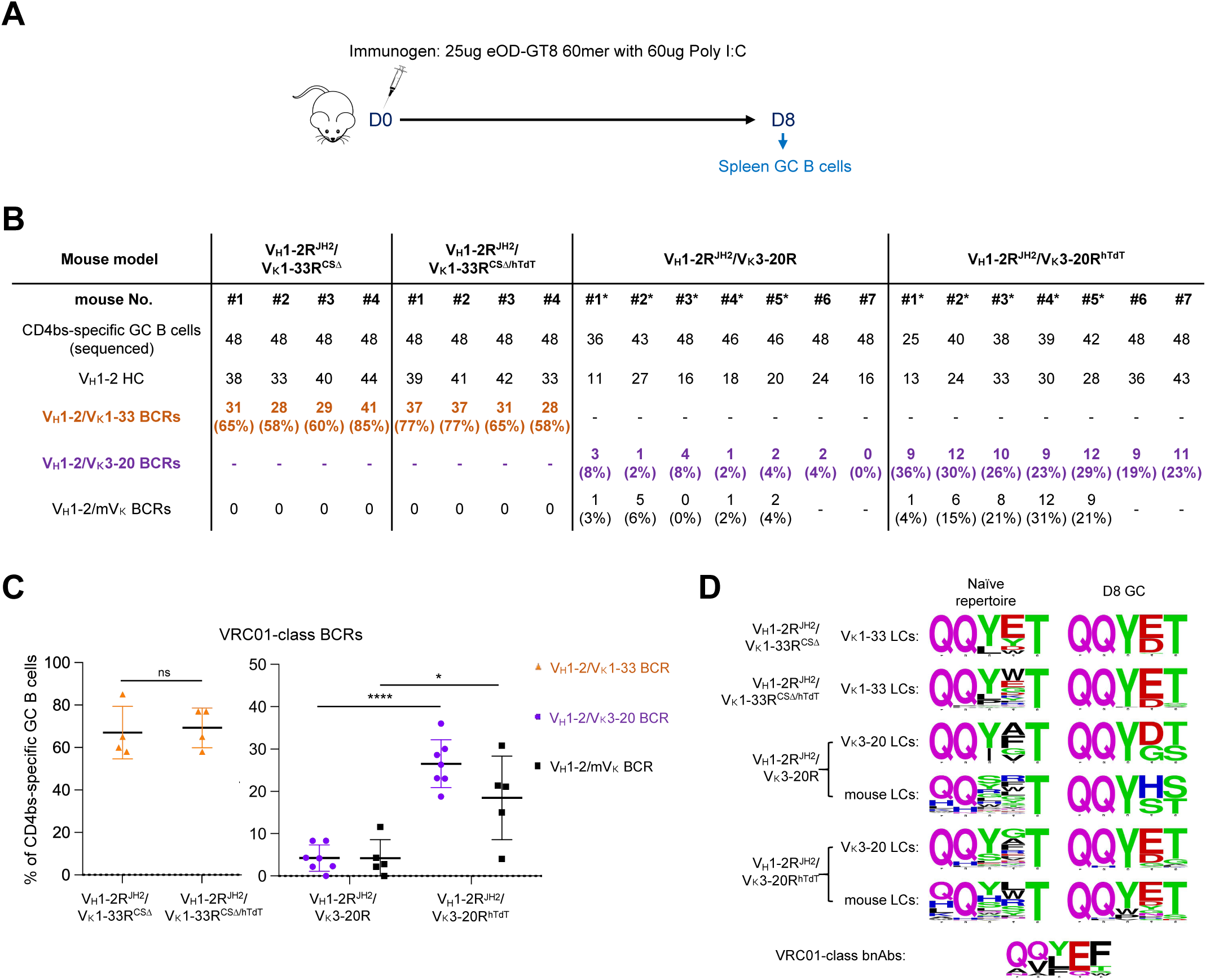
Enforced hTdT expression enhances the VRC01-class GC responses induced by eOD-GT8 60mer. (A) Immunization scheme (see text for details). (B) Summary of all VRC01-class BCR sequence information obtained from eOD-GT8 immunization in V_H_1-2R^JH2^/Vκ1-33R^CSΔ^, V_H_1-2R^JH2^/Vκ1-33R^CSΔ/hTdT^, V_H_1-2R^JH2^/Vκ3-20R and V_H_1-2R^JH2^/Vκ3-20R^hTdT^ mice. VRC01-class BCRs were defined by V_H_1-2 HCs pairing with Vκ1-33/Vκ3-20/mouse LCs with 5-aa CDR3s. Statistical analyses are shown in (C). * indicates the mouse harboring a mutated Vκ3-20 allele. (C) The frequency of VRC01-class BCRs expressed in CD4bs-specific GC B cells from V_H_1-2R^JH2^/Vκ1-33R^CSΔ^, V_H_1-2R^JH2^/Vκ1-33R^CSΔ/hTdT^, V_H_1-2R^JH2^/Vκ3-20R and V_H_1-2R^JH2^/Vκ3-20R^hTdT^ mice. Each point represents one mouse. *p* values were calculated by unpaired, two-tail t-test. \**p* <0.05, \*\**p* <0.01, \*\*\**p* <0.001, \*\*\*\**p* <0.0001 (D) 5-aa LC CDR3 sequence logos for Vκ1-33, Vκ3-20 and mouse LCs in naïve BCRs (left column) and eOD-GT8 60mer-induced VRC01-class BCRs at day 8 post-immunization (right column). The sequences of 5- aa LC CDR3s in naïve B cells were derived from HTGTS-rep-seq data shown in Fig.1B, Fig.S1, E and F, and Fig. S4A. The sequences of 5-aa LC CDRs in eOD-GT8 60mer-induced VRC01-class BCRs were recovered from V_H_1-2R^JH2^/Vκ1-33R^CSΔ^, V_H_1-2R^JH2^/Vκ1-33R^CSΔ/hTdT^, V_H_1-2R^JH2^/Vκ3-20R and V_H_1-2R^JH2^/Vκ3-20R^hTdT^ mice shown in (B). For comparison, the 5-aa LC CDR3 sequences for VRC01-class bnAbs were shown in bottom.

On day 8 post-immunization, the Glu96 (E), a conserved residue in 5-aa LC CDR3s of VRC01-class bnAbs, was dominantly selected by eOD-GT8 in VRC01-class 5-aa LC CDR3s from V_H_1-2R^JH2^/Vκ1-33R^CSΔ^, V_H_1-2R^JH2^/Vκ1-33R^CSΔ/hTdT^ and V_H_1-2R^JH2^/Vκ3-20R^hTdT^ mice, but not from V_H_1-2R^JH2^/Vκ3-20R mice (Fig.2D; Fig.S7D). This finding indicated that Vκ to Jκ joining events involving Vκ3-20 or mouse Vκs in the Vκ3-20 mice require N regions added by hTdT to generate the critical E residue in the VRC01-class 5-aa CDR3. Examination of Vκ3-20 and mouse Vκ sequences proved that this is the case (Fig.S7E). On the other hand, examination of the Vκ1-33 sequences confirm that they can directly form the E residue in the VRC01-class 5-aa CDR3 when joined to mouse Jκ1 and human Jκ1 in the absence of hTdT activity (Fig.S7E). Lack of this E residue in 5-aa mouse LC CDR3s in primary GCs that arose after a single eOD-GT8 immunization was also noted in prior studies (46, 49, 63, 64). Thus, hTdT expression substantially enhanced the VRC01/Vκ3-20 and VRC01/mVκ GC response to eOD-GT8 immunization by generating Vκ3-20-based VRC01-class 5-aa CDR3s that, as a result of N-region addition, have the capacity to encode the critical CDR3 E residue.

### Generation of V_H_1-2^JH2^/Vκ1-33/Vκ3-20^hTdT^-rearranging mice

We bred the V_H_1-2R^JH2^/Vκ1-33R^CSΔ/hTdT^, V_H_1-2R^JH2^/Vκ3-20R^hTdT^ mouse lines together to make an even more human-like model that rearranges both VRC01-class Vκs. In this new V_H_1-2R^JH2^/Vκ1-33R^CSΔ/hTdT^/Vκ3-20R^hTdT^ mouse model, Vκ1-33 and Vκ3-20 LCs were expressed in 7.8% and 3.4% of splenic B cells, respectively (Fig.S8A). However, on day 8 post-immunization with eOD-GT8 60mer, VRC01/Vκ3-20 GC B cells were outcompeted by VRC01/Vκ1-33 GC B cells and were hardly represented in GCs, suggesting the frequency or affinity of responding VRC01/Vκ1-33 precursors was much higher than that of VRC01/Vκ3-20 precursors in this model (Fig.S8B). Thus, we further generated V_H_1-2R^JH2^/Vκ1-33R/Vκ3-20R^hTdT^ model, in which Cer/Sis is still present on the Vκ1-33 allele, leading to a reduction in Vκ1-33 LC-expressing splenic B cell frequency to 0.74% (Fig.3, A and B). Indeed, the relative frequency of Vκ1-33 versus Vκ3-20 expressing splenic B cells in the V_H_1-2R^JH2^/Vκ1-33R/Vκ3-20R^hTdT^ model are more comparable to that of humans (65). To assess the frequency of VRC01-precursors, we sorted eOD-GT8-specific naïve B cells and identified their BCR sequences (Fig. 3C and Fig. S8C). The frequency of eOD-GT8-specific VRC01 precursors using Vκ1-33 or Vκ3-20 LCs in this mouse model was approximately 1 in 230,000 (VRC01/Vκ1-33: 1 in 500,000; VRC01/Vκ3-20: 1 in 420,000) (Fig. 3D), which is comparable to approximately 1 in 400,000 frequency of eOD-GT8-specific VRC01 precursors measured in humans (44). We also estimated the VRC01-precursor based on HTGTS-Rep-seq data by multiplying the frequency of V_H_1-2 HCs by the frequency of Vκ1-33 and Vκ3-20 LCs with 5-aa CDR3s (Fig. 3E). The results suggest that only a small proportion of B cells expressing V_H_1-2 HCs and Vκ3-20 LCs with 5-aa CDR3s bound to eOD-GT8.

**Figure 3.**
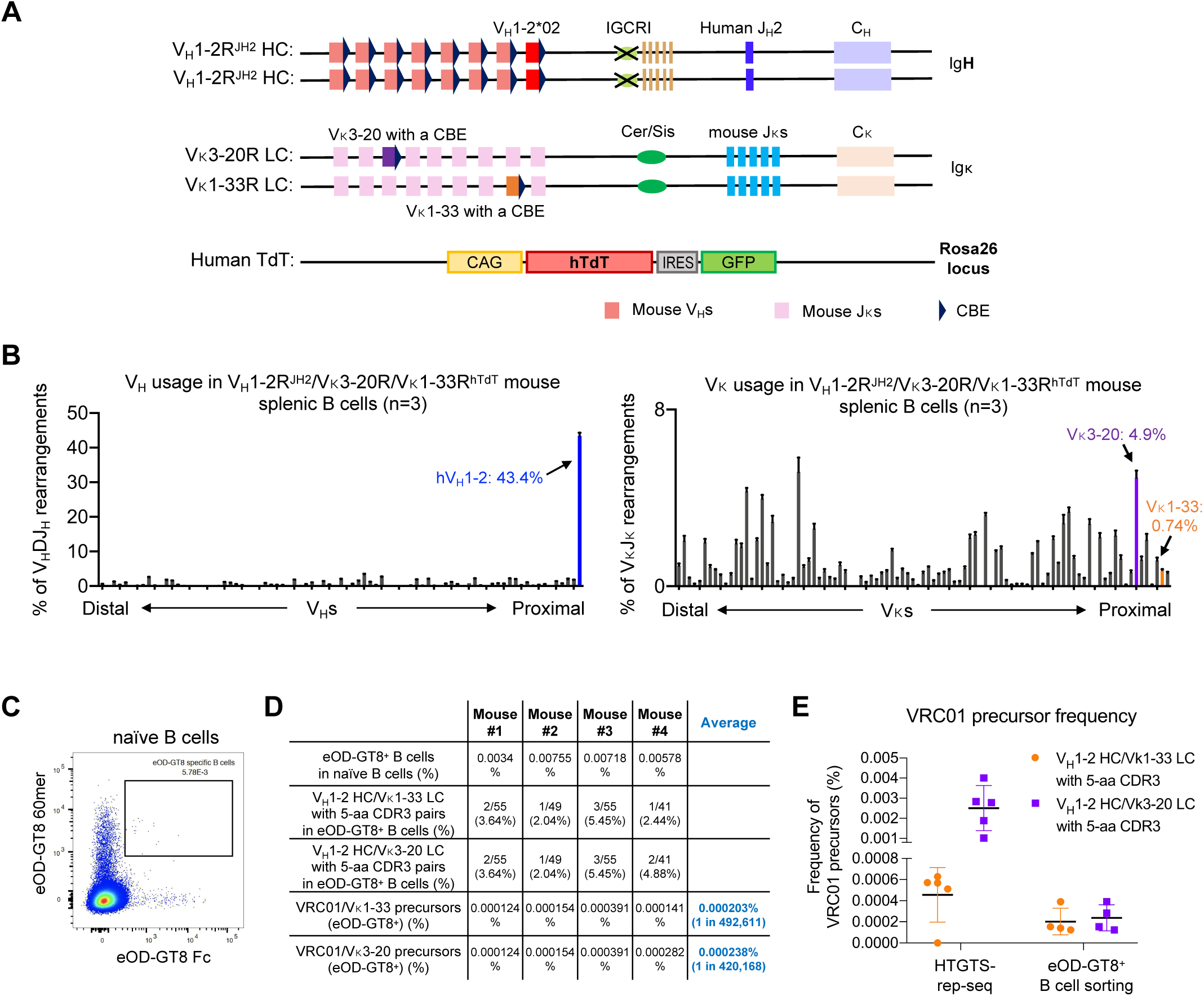
Generation and characterization of the V_H_1-2^JH2^/Vκ1-33/Vκ3-20^hTdT^-rearranging mouse models. (A) Illustration of genetic modifications in the IgH and Igκ locus of V_H_1-2^JH2^/Vκ1-33/Vκ3-20^hTdT^-rearranging mouse models. The *Jκ*-proximal Vκ3-2 was replaced with human Vκ1-33 plus a CBE 50bp downstream of its RSS. (B) HTGTS-rep-seq analysis of V_H_ (upper panel) and Vκ (bottom panel) usage in V_H_1-2^JH2^/Vκ1-33/Vκ3-20^hTdT^-rearranging mouse splenic B cells. The x axis listed all functional V_H_s or Vκs from the distal to the *D-* or *Jκ*-proximal end. The histogram displayed the percent usage of each V_H_ or Vκs among all productive V_H_DJ_H_ or VκJκ rearrangements. The usage of human V_H_1-2, Vκ1-33 and Vκ3-20 were shown in blue, orange and purple, respectively. Data from (A) and (B) were mean ± SD of five libraries from different mice. (C) FACS analyses of eOD-GT8-specific naïve B cells in V_H_1-2^JH2^/Vκ1-33/Vκ3-20^hTdT^-rearranging mouse. The boxed eOD-GT8-specific naïve B cells were sorted for single cell sequencing. (D) Summary of VRC01 precursor sequence information obtained from naïve B cell repertoire. The eOD-GT8 specific B cells were defined by eOD-GT8 60mer^+^ and eOD-GT8 Fc^+^. The final frequency of VRC01 precursors in V_H_1-2^JH2^/Vκ1-33/Vκ3-20^hTdT^-rearranging mouse models is 1 in 226,757, approximately. (E) Frequency of VRC01 precursors in V_H_1-2^JH2^/Vκ1-33/Vκ3-20^hTdT^-rearranging mice measured by HTGTS-rep-seq or eOD-GT8-specific B cell sorting. The VRC01 precursors were defined by V_H_1-2 HCs pairing with Vκ1-33 and Vκ3-20 LCs with 5-aa CDR3s.

### VRC01-class B cells develop SHM and affinity maturation in GCs induced by eOD-GT8 60mer

To test if V_H_1-2R^JH2^/Vκ1-33R/Vκ3-20R^hTdT^ mice respond to the VRC01-class prime immunogens and support affinity maturation of VRC01-class GC B cells at sufficient levels to support future prime-boost studies, we immunized them with eOD-GT8 60mer and then boosted them with eOD-GT8 60mer at day 28 (Fig.4A). VRC01/Vκ1-33, VRC01/Vκ3-20 and VRC01/mVκ B cells were highly enriched in CD4bs-specific GC B cells at both 8 day and 36 day post-immunization (Fig.4B; Fig.S9, A to C). Evaluation of GC responses at day 8 and day 36 revealed that the frequencies of VRC01/Vκ3-20 GC B cells and VRC01/Vκ1-33 GC B cells were comparable at day 8, but the frequencies of VRC01/Vκ3-20 GC B cells was higher than that of VRC01/Vκ1-33 GC B cells at day 36 (Fig.4C). Sequencing analyses of VRC01-class antibodies cloned from both day 8 and day 36 GCs revealed extensive SHM, with a maximum of 17 aa mutations and a median of 9 aa mutations at day 36 (Fig.4D; Fig.S9, D and E), and wide ranges of HC CDR3 length (Fig. S9F). To further analyze VRC01-class GC B cell sequence mutations, we compared them to intrinsic mutation patterns generated from non-productive rearrangements of GC B cells without affinity selection (Fig.4E; Fig.S9, G to I) (see Method) (66). The Q61R mutant on the V_H_1-2 HC reported for VRC01-class bnAbs was significantly enriched in day 36 VRC01-class antibodies (Fig.4F) (42). The Glu96 (E) residues in LC CDR3s were dominant in all types of day 36 VRC01-class antibodies (Fig.4G). We expressed several VRC01-class antibodies with different LCs cloned from day 8 and day 36 GCs. Antibodies from day 8 GCs showed a range of binding affinities, with a median of 100nM K_D_, to eOD-GT8 (Fig.4H). For the antibodies from day 36 GCs, about 50% showed much higher binding activities, below 1nM K_D_, representing an average affinity improvement of 100-fold (Fig.4H and Table S1). Altogether, our findings strongly indicate that the V_H_1-2R^JH2^/Vκ1-33R/Vκ3-20R^hTdT^ VRC01-class and related models will facilitate testing prime-boost immunization strategies aimed to advance eOD-GT8-primed vaccination studies to be used in human clinical trials.

**Figure 4.**
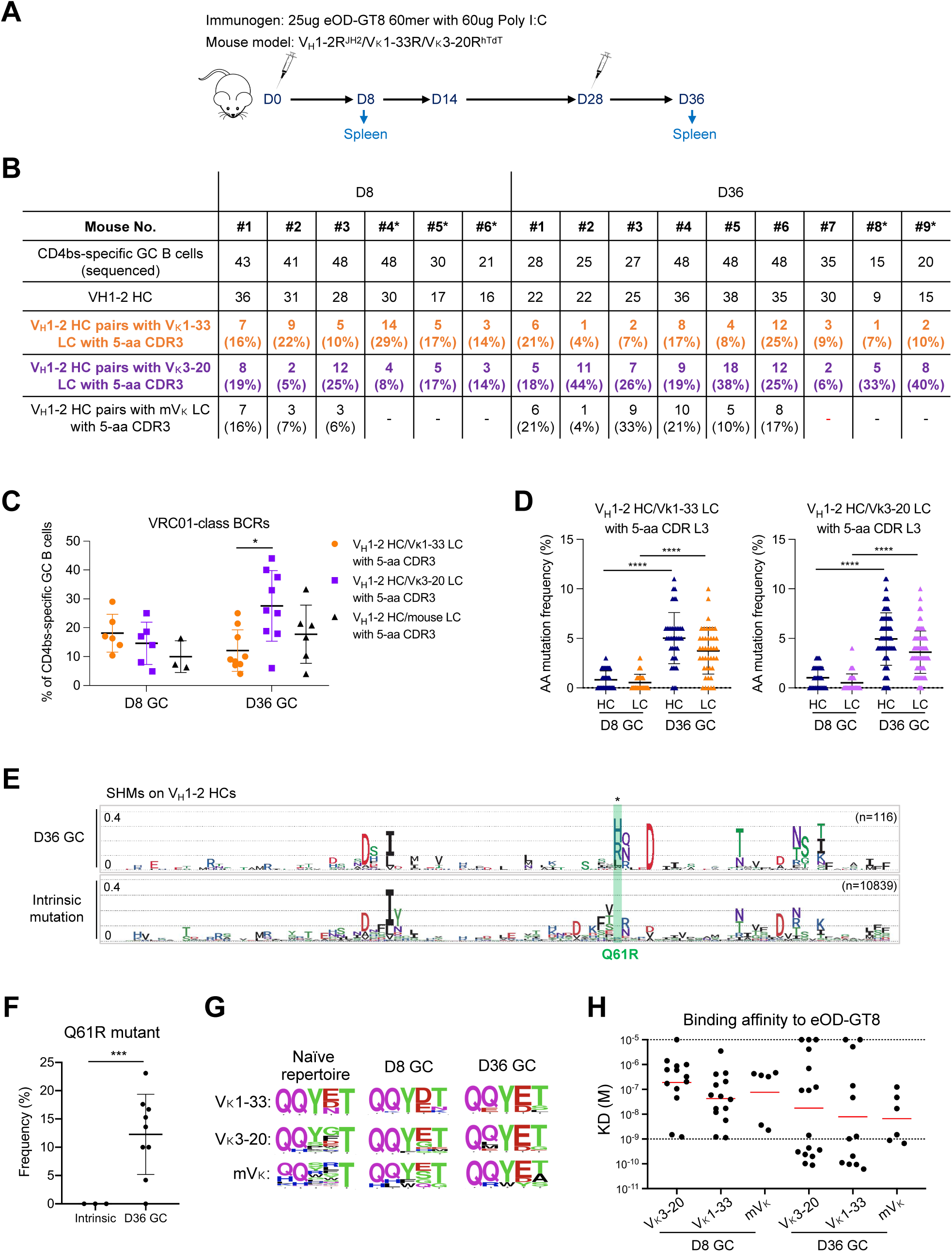
Strong VRC01-class GC responses induced by eOD-GT8 60mer in V_H_1-2^JH2^/Vκ1-33/Vκ3-20^hTdT^-rearranging mouse models. (A) Immunization scheme (see text for details). (B) Summary of all VRC01-class BCR sequence information obtained from eOD-GT8 60mer immunization at day 8 and day 36. VRC01-class BCRs were defined by V_H_1-2 HCs pairing with Vκ1-33/Vκ3-20/mouse LCs with 5-aa CDR3s. Statistical analyses are shown in (C). * indicates the mouse harboring a mutated Vκ3-20 allele. (C) The frequency of VRC01-class BCRs expressed in CD4bs-specific GC B cells from day 8 and day 36 GCs of V_H_1-2R^JH2^/Vκ1-33R/Vκ3-20R^hTdT^ mice. (D) Amino acid mutation frequency in VRC01-class antibodies cloned from day 8 and day 36 GCs of V_H_1-2R^JH2^/Vκ1-33R/Vκ3-20R^hTdT^ mice. Each dot represents one HC or one LC. The median with interquartile range is plotted. (E) Mutation frequency of each amino acid on germline-encoded V_H_1-2 region of VRC01-class antibodies cloned from day 36 GCs shown in sequence logo profiles. For reference, the intrinsic mutation patterns from non-productive rearrangements are represented on the bottom (see method for details). The distance between dotted horizontal lines representing 0.1 (10%). The Q61R mutant is labeled in green. (F) Frequency of Q61R mutant on day 36 V_H_1-2 HC compared to that in intrinsic mutation patterns. (G) 5-aa LC CDR3 sequences in naïve repertoire and VRC01-class antibodies cloned from day 8 and day 36 GCs induced by eOD-GT8 60mer. 5-aa LC CDR3 sequence logos for Vκ1-33, Vκ3-20 and mouse LCs in naïve BCRs (left column), 8-day GCs (middle column) and 36-day GCs (right column) induced by eOD-GT8 60mer. (H) eOD-GT8 dissociation constants measured by surface plasmon resonance (SPR) for eOD-GT8 60mer elicited VRC01-class antibodies (see method and Supplementary Table for details). Data are shown for VRC01-class antibodies from 8-day and 36-day GCs. Bars represent geometric mean (red). Statistical comparisons in (**C**), (**D**) and (**F**) were performed using a two-tailed unpaired t test. \**p* <0.05, \*\**p* <0.01, \*\*\**p* <0.001, \*\*\*\**p* <0.0001

## Discussion

Many prior mouse models employed to test vaccine strategies designed to elicit VRC01-class HIV-1 bnAbs had exceedingly high or extremely low levels of VRC01-class precursor B cells. Other approaches to generate more physiological levels of VRC01 precursors in mouse models were limited by being designed to test only the eOD-GT8 priming immunogen in the context of very limited precursor diversity. We have now described more physiologically relevant VRC01-class V(D)J-rearranging mouse models for testing priming and boosting strategies designed to elicit VRC01-class bnAbs. These new VRC01-class rearranging mouse models rearrange both human VRC01-class V_H_1-2 and Vκ3-20 and/or Vκ1-33 variable region gene segments, along with mouse V_H_s and Vκs during normal B cell development. The various mouse lines generated to make the VRC01-class rearranging models described here employ several different genetic strategies that should allow titration of the expression level of diverse Vκ3-20- and/or Vκ1-33-based variable region exons to establish mouse models that generate VRC01 precursor B cells over a wide range of levels (Table S2). Of these models the V_H_1-2R^JH2^/Vκ1-33R/Vκ3-20R^hTdT^ model, described in depth in this report, generates a highly diverse set of potential VRC01-class precursors in mouse repertoires at similar relative levels to those found in human B cell repertories. Importantly, the potential VRC01-class precursors with highly diverse CDR3s generated in the VRC01-class rearranging models should not be biased with respect to evaluating the efficacy of any particular VRC01-class priming immunogen (Fig. S9F).

In this initial study, we have tested the eOD-GT8 priming immunogen in several VRC01 class rearranging models, including the most human-like V_H_1-2R^JH2^/Vκ1-33R/Vκ3-20R^hTdT^ model, and found robust engagement of VRC01-class precursors into GCs where they generated equally robust eOD-GT8-specific responses. Other types of priming immunogens that may not be as robust in engaging VRC01-class precursors as eOD-GT8, such as 426c-degly3 Ferritin (40, 47) or GT1 trimer (67), should also be able to be readily evaluated in our new models. Conceivably, studies of some VRC01-class immunogens that have lower affinity for precursors may benefit initially through the use of VRC01-class models that express higher levels of VRC01-class precursors (Table S2). Also, as individual VRC01-class precursor B cells in these new VRC01-class rearranging models express one of a multitude of different variations of the potential VRC01 precursor, they may, in theory, be useful for identifying new pathways that could lead to the generation of potent VRC01-class bnAbs. For any tested priming immunogen that generates a response, our new models could also be used to test sequential boost immunogens designed to lead them through rounds of SHM/affinity maturation that drive responses towards the generation of VRC01-class bnAbs as described for less diverse earlier versions of these models (49, 68).

A key feature of our new models is their ectopic TdT expression that forces their mouse Pre-B cells to further diversify their mouse and human LC variable region repertoires and make them more human-like both with respect to contributing N-region diversity and by dampening recurrent MH-mediated join levels in their postnatal LC repertoires. As mentioned, absence of TdT in fetal repertoires promotes recurrent MH-mediated junctions that lead to generation of particular Ig or TCR variable region exon sequences (14–21). For example, generation of recurrent “canonical” joins in fetal repertoires in the absence of TdT and N region additions underlies generation of canonical junctions encoding recurrent γ*/*δ TCRs expressed on “innate-like” intraepithelial γ*/*δ T cells that persist into adulthood in both mice and humans (69, 70). Notably, enforced TdT expression during fetal lymphocyte development dampens some such responses (13, 28). In this study, we found that enforced TDT expression in mouse Pre-B cells increased the frequency of short 5-aa CDR3 sequences, such as those used in a VRC01-class response, and promoted a specific Vκ3-20-based eOD-GT8 primary response by generating N sequences that contribute to encoding a critical VRC01 class 5-aa CDR3 residue. Analyses of human Vκ3-20-based VRC01-class sequences indicate that this mechanism also operates in humans (e.g. Fig. S7E). By extension, it is likely that postnatal TdT expression will similarly contributes to other responses.

The strategies we employed for constructing the VRC01 rearranging mouse model can be generally adopted for generating mouse models for other classes of anti-HIV-1 bnAbs. In this regard, CDR3 diversification, including engineering the models to make very long human CDR3s, will be especially relevant for testing immunogens for bnAbs that rely heavily on CDR3 to contact Env epitopes, such as those of the V2 apex, V3 glycan and MPER classes (71). The limitations with previously employed strategies to generate mouse models to test VRC01-class immunization strategies outline above also will apply to mouse models designed to test immunogens in the context of these other bnAb lineages. Beyond this, all straight precursor variable region knock-in strategies are limited by difficulty in accurately inferring the CDR3 of the common unmutated ancestor sequence of precursors, which may include contributions from both non-templated nucleotides and somatic hypermutations (72). Second, due to the enormous CDR3 diversity in human antibody repertoires, a specific bnAb precursor may not be present in all individuals. To work at a population level, a vaccine should stimulate B cells expressing a range of related precursors. Mouse models expressing a unique bnAb precursor cannot assess this critical parameter. Also, expression of certain bnAb precursors HCs or LCs can interfere with B cell development, leading to B cell deletion in bone marrow or anergy in peripheral lymphoid tissues (71, 73–75). The prototype VRC01-class rearranging mouse model we have described here, addresses these potential issues in the VRC01 lineage, as V(D)J recombination generates human VRC01-class precursors that express highly diverse CDR3s, many of which may be compatible with bnAb development. Thus, this type of mouse HIV-1 vaccine model does not depend on UCA inference. Additionally, the CDR3 diversity in the model facilitates assessment of the ability of immunogens to tolerate CDR3 flexibility and mobilize related precursors for bnAb development. Finally, by generating diverse human primary BCR repertoires, rearranging mouse models can provide precursors that support normal B cell development and, correspondingly, generate B cells responsive to immunization.

## Materials and Methods

### VRC01-rearranging mouse model and embryonic stem cells

The genetic modifications in the *Igκ* locus were introduced into previously generated V_H_1-2 ES cells (129/Sv and C57BL/6 F1 hybrid background), using targeting strategies described previously (49). The mouse Vκ3-7 segment was replaced with human Vκ3-20 segment with an attached CTCF-binding element (CBE) (<u>atccaggaccagcagggggcgcggagagcacaca</u>) inserted 50 bp downstream of human Vκ3-20 segment. The replacement was mediated by homologous recombination using a PGKneolox2DTA.2 (Addgene #13449) construct and one guide RNA that targeted the mouse Vκ3-7 segment. The human TdT cDNA was cloned into CTV (Addgene #15912) construct in which the TdT expression was driven by CAG promotor and followed by a EGFP expression that mediated by an internal ribosome entry site (IRES) (76). The TdT expression cassette was inserted into the first intron of mouse Rosa26 gene which is on the same chromosome 6 with *Igκ* locus by homologous recombination. The sequence of guide RNA used for targeting were listed in Table S3. The ESCs were grown on a monolayer of mitotically inactivated mouse embryonic fibroblasts (iMEF) in DMEM medium supplemented with 15% bovine serum, 20mM HEPES, 1x MEM nonessential amino acids, 2mM Glutamine, 100 units of Penicillin/Streptomycin, 100 mM b-mercaptoethanol, 500 units/ml Leukemia Inhibitory Factor (LIF).

The V_H_1-2^JH2^/Vκ3-20^hTdT^ -rearranging mouse was generated by blastocyst injection of the ES cells described above and several rounds of breeding to get germline transmission and homozygous mice. The V_H_1-2^JH2^/Vκ1-33/Vκ3-20^hTdT^-rearranging mouse was generated by cross breeding of V_H_1-2^JH2^/Vκ3-20^hTdT^ and V_H_1-2^JH2^/Vκ1-33^hTdT^ mice. Thus, human Vκ1-33 and Vκ3-20 segments were used on separated alleles. All mouse experiments were performed under protocol 20-08-4242R approved by the Institutional Animal Care and Use Committee of Boston Children’s Hospital.

### Immunogen and Immunization

Immunogen eOD-GT8 60mer was made as previously described (49). For immunization, each 8-12 weeks old mouse was immunized with 200 ul mixture that contain 25 ug filter-sterilized immunogen and 60 ug of poly I:C in PBS by intraperitoneal injection.

### Splenic B cell, GC B cell purification and Antigen-specific GC B Cell Sorting

Splenic B cells used for HTGTS-Rep-seq were purified from unimmunized 5-8 weeks old mice by MACS® Microbeads according to the manufacturer’s protocol. In brief, spleens were dissected out from unimmunized mice, prepared into single cell suspensions and stained with anti-B220 Microbeads for 20 minutes at 4°C. The splenic B cells were collected using the LS column and MACS^TM^ Separator. GC B cells used for Rep-SHM-seq were purified from 8-12 weeks old mice after eOD-GT8 60mer immunization. GC B cells were sorted for the phenotype B220^+^ (BV711: BioLegend 103255), CD95^+^ (PE-Cy7: eBioscience 557653) and GL7^+^ (PE: BioLegend 144607). CD4-binding site-specific GC B cells for single cell RT-PCR were further selected for the phenotype eOD-GT8 Fc^+^ and ΔeOD-GT8 Fc^-^. The eOD-GT8 Fc was conjugated with Alexa Fluor 647 fluorescence (Thermo Fisher Scientific A30009). The ΔeOD-GT8 Fc was conjugated with Biotin (Thermo Fisher Scientific A30010) and then stained with BV605 (BioLegend 405229).

### Human tonsil mature naïve B cell isolation and genomic DNA extraction

Human tonsils were obtained from discarded tissues as part of a routine tonsillectomy from patients at Boston Children’s Hospital. Human tissues were obtained under the IRB approved protocol IRB-P00026526, to J.P.M. Tonsils were minced in RPMI 1640 with 10% FBS and forced through a 45 μm mesh and washed twice with media. The single cell suspension was stained with 7-AAD (Biolegend) for viability and antibodies directed against human CD19 (APC clone SJ25-C1, Thermo Fisher Scientific), CD38 (PE-Cy7 clone HB-7, Biolegend), IgD (FITC polyclonal, Thermo fisher) and CD27 (APC-Cy7 clone M-T271, Biolegend). Live Naïve B cells were obtained by sorting the stained cells using a FACS (fluorescence-activated cell sorting) Aria (BD Biosciences) as 7-AAD^-^CD19^+^CD38^-^IgD^+^CD27^-^. Genomic DNA from sorted cells was prepared using a DNeasy Blood and Tissue Kit (Qiagen) according to the manufacturer’s protocol.

### HTGTS-Rep-seq and Rep-SHM-seq Analysis

10 ug of DNA from purified splenic B cells was used for generating HTGTS-Rep-seq libraries as previously described (77). 4 bait primers that target mouse Jκ1, Jκ2, Jκ4 and Jκ5 were mixed to capture all Igκ light chain repertoire in one library. One bait primer that targets human J_H_2 was used to capture heavy chain repertoire. The sequences of human J_H_2 and mouse Jκ primers were as same as the previously reported (54, 66). These HTGTS-Rep-seq libraries were sequenced by Illumina NextSeq 2 x 150-bp paired end kit analyzed with the HTGTS-Rep-seq pipeline (77). DNA from GC B cells was used for generating Rep-SHM-seq libraries as previously described (66). To capture the full-length V(D)J sequence especially the CDR1 region for intrinsic SHM analysis, we designed bait primers that target human V intron regions. The primer sequences are in Table S3. These Rep-SHM-seq libraries were sequenced by Illumina MiSeq 2 x 300-bp paired end kit analyzed with the Rep-SHM-seq pipeline, which uses IgBLAST to annotate V, D, J and CDRs for each read (66).

### Single Cell RT-PCR and monoclonal antibody production

Single cell RT-PCR were performed as described previously (57). In brief, single antigen-specific GC B cells were sorted into 96-well plate that contain 5ul of lysis buffer in each well. After sorting, we used primers mixture that specifically target Cµ, Cγ1, Cγ2a and Cκ to perform reverse transcription and then two rounds of nested PCR to amplify the V(D)J sequences of V_H_1-2 heavy chain, mouse light chain, human Vκ3-20 and Vκ1-33 light chain. PCR products were run on agarose gels and perform sanger sequencing to confirm their identity. The primer sequences for V_H_1-2 HC, Vκ3-20 and Vκ1-33 LC amplification were in Table S3. The primer sequences for mouse LC amplification were as same as previously reported (78). The antibody expression constructs containing the heavy-chain and the light-chain variable region exons, with human constant region sequences (IgG1, Igκ) at the C terminus were made by Genscript. Monoclonal antibodies were generated using the Expi293 expression system (Thermo Fisher Scientific) and purified by high-performance liquid chromatography (HPLC) coupled with HiTrap Protein A HP columns (Cytiva).

### Carterra Human IgG Capture

Kinetics and affinity of antibody-antigen interactions were measured on Carterra LSA using HC30M or CMDP Sensor Chip (Carterra) and 1x HBS-EP+ pH 7.4 running buffer (20x stock from Teknova, Cat. No H8022) supplemented with BSA at 1mg/ml. Chip surfaces were prepared for ligand capture following Carterra software instructions. In a typical experiment about 1000-1700 RU of capture antibody (SouthernBiotech Cat no 2047-01) in 10 mM Sodium Acetate pH 4.5 was amine coupled. Phosphoric Acid 1.7% was our regeneration solution with 30 seconds contact time and injected three times per each cycle. Solution concentration of ligands was above 10ug/ml and contact time was 10min. as per Carterra manual. Raw sensograms were analyzed using Kinetics software (Carterra), interspot and blank double referencing, Langmuir model. Analyte concentrations were quantified on NanoDrop 2000c Spectrophotometer using Absorption signal at 280 nm.

### Analyses of CDR3 diversity and MH-mediated V(D)J recombination

The lengths of insertion and MH for Vκ to Jκ rearrangement were annotated based on HTGTS-Rep-seq results. Insertion nucleotides can be classified into P (palindromic) nucleotides and N (non-template) nucleotides. For a read that can be aligned to the 3’ end of V segment or 5’ end of J segment, the length of P nucleotides was determined by greedy alignment of read sequence outside the V or J end to the reverse complimentary V or J sequence from the end. And the remaining insertion nucleotides were classified as N nucleotides. The length of MH was determined by the length of overlapping read sequence that could be aligned to both V and J (V_end_on_read – J_start_on_read + 1) after greedy alignment to V and J. CDR3 diversity was represented by the percentage of unique CDR3s for a series of downsampled read numbers (e.g. 20, 50, 100, 200), which could be viewed as rarefaction and estimated by R package ‘iNEXT’. Welch’s t-test was used to compare the percentage of unique CDR3s between groups.

### Statistical analysis

Statistical tests with appropriate underlying assumptions on data distribution and variance characteristics were used. t-test was used as indicated in the figure legends. Statistical analysis was performed in Prism (v.8, GraphPad Software).

### Data and software availability

All data needed to evaluate the conclusions of the paper are presented in the paper or deposited on the online database. Nucleotide sequences have been deposited to GenBank (accession Nos. OP598882 - OP599353). The next-generation sequencing data reported in this paper have been deposited in the Gene Expression Omnibus (GEO) database under the accession number GSE214884. The computational pipeline of Rep-SHM-Seq and the code for statistical analysis tools used in this study are available at https://github.com/Yyx2626/HTGTSrep

## Acknowledgments

We thank the F.W.A. laboratory members for discussions and comments, Hwei-Ling Cheng for the advice about ES cell culture and Jianqiao Hu for the bioinformatics assistance. We thank Tina-Marie Mullen for antibody production. This work was supported by the Bill & Melinda Gates Foundation INV-021989 (F.W.A.) and R01 AI100887 (J.P.M.). F.W.A. is an Investigator of the Howard Hughes Medical Institute.

---

**Figure S1.**
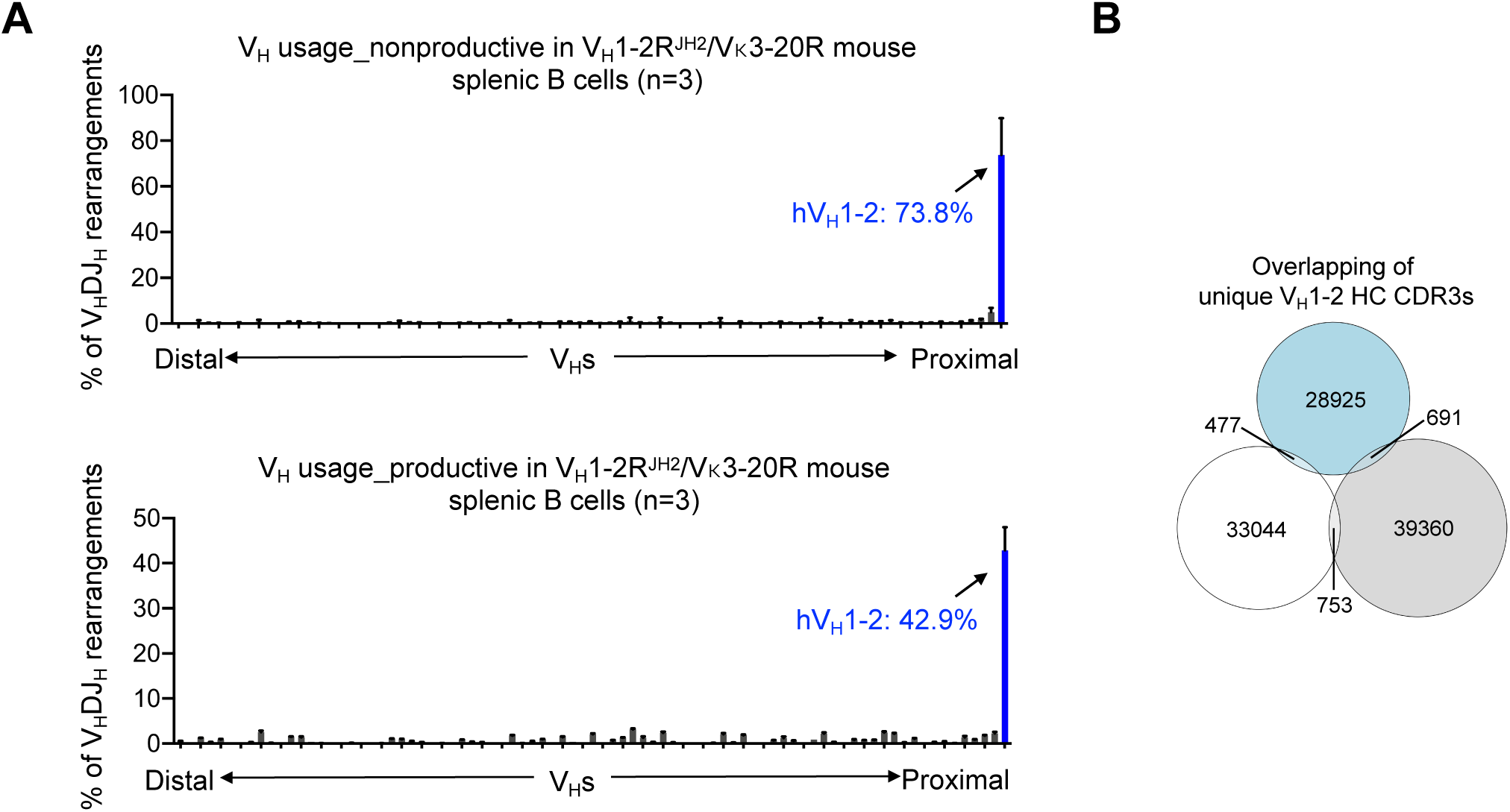
Characterization of V_H_1-2^JH2^-rearranging heavy chain. (A) HTGTS-rep-seq analyses of V_H_ non-productive (upper panel) and productive (bottom panel) rearrangements in V_H_1-2^JH2^/Vκ3-20-rearranging splenic B cells. The histogram displays the percent of nonproductive or productive rearrangements of each V_H_ among all V_H_DJ_H_ nonproductive or productive rearrangements. The frequencies of V_H_ nonproductive rearrangements represent the V_H_ usages in primary V(D)J rearrangements, as the nonproductive allele was not under selection during B cell development. Data were average of 3 experimental repeats with error bars representing SDs. (B) Venn diagram showed the V_H_1-2 HC CDR3 diversity. The unique reads derived from the same libraries in Fig. 1B.

**Figure S2.**
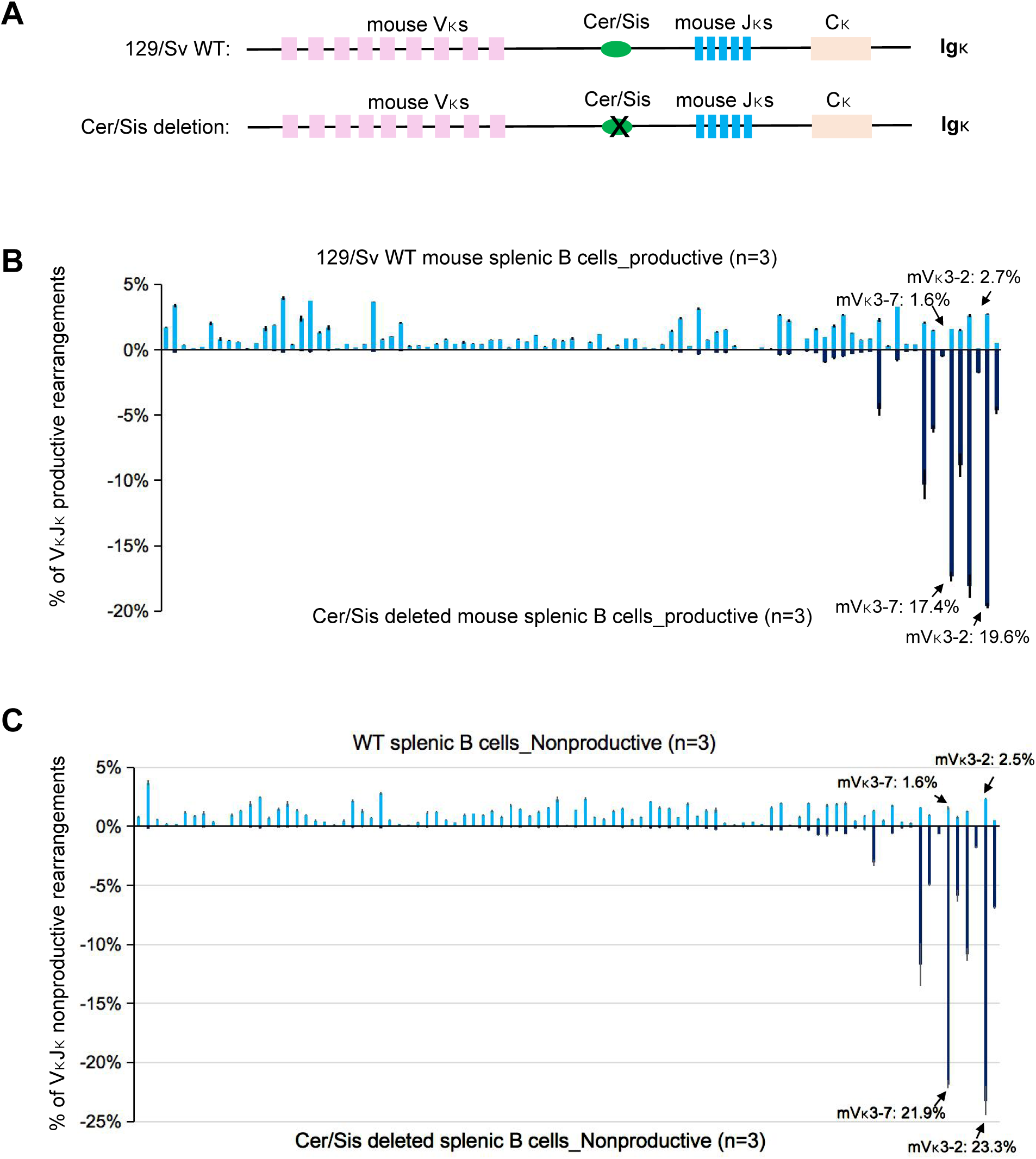
Cer/sis deletion in wild-type mice increased the utilizations of proximal Vκs, including Vκ3-2 and Vκ3-7. (A) Illustration of Cer/sis deletion in the Igκ locus. The strategy of Cer/sis deletion was the same as recently described (35). (B) HTGTS-rep-seq analyses of Vκ usages in wild type (upper panel) and Cer/Sis deleted (lower panel) mouse splenic B cells. The x axis lists all functional Vκs from the distal to the *Jκ*-proximal ends. The histogram displays the percent usage of each Vκ among all productive VκJκ rearrangements. The productive Vκ rearrangements in splenic B cells represent the Vκ usage in the naïve BCR repertoire. The data in wild type mouse splenic B cells were derived from our recent study (35). (C) HTGTS-rep-seq analyses of Vκ nonproductive rearrangements in wild type (upper panel) and Cer/sis deleted (bottom panel) splenic B cells. The histogram displays the percent of nonproductive rearrangements of each Vκ among all nonproductive VκJκ rearrangements. The percentage of Vκ segments in nonproductive rearrangements represents the V usage in primary V(D)J recombination. The data in wild type mouse splenic B cells were derived from our recent study (35). Data from (B) and (C) were average of 3 experimental repeats with error bars representing SDs.

**Figure S3.**
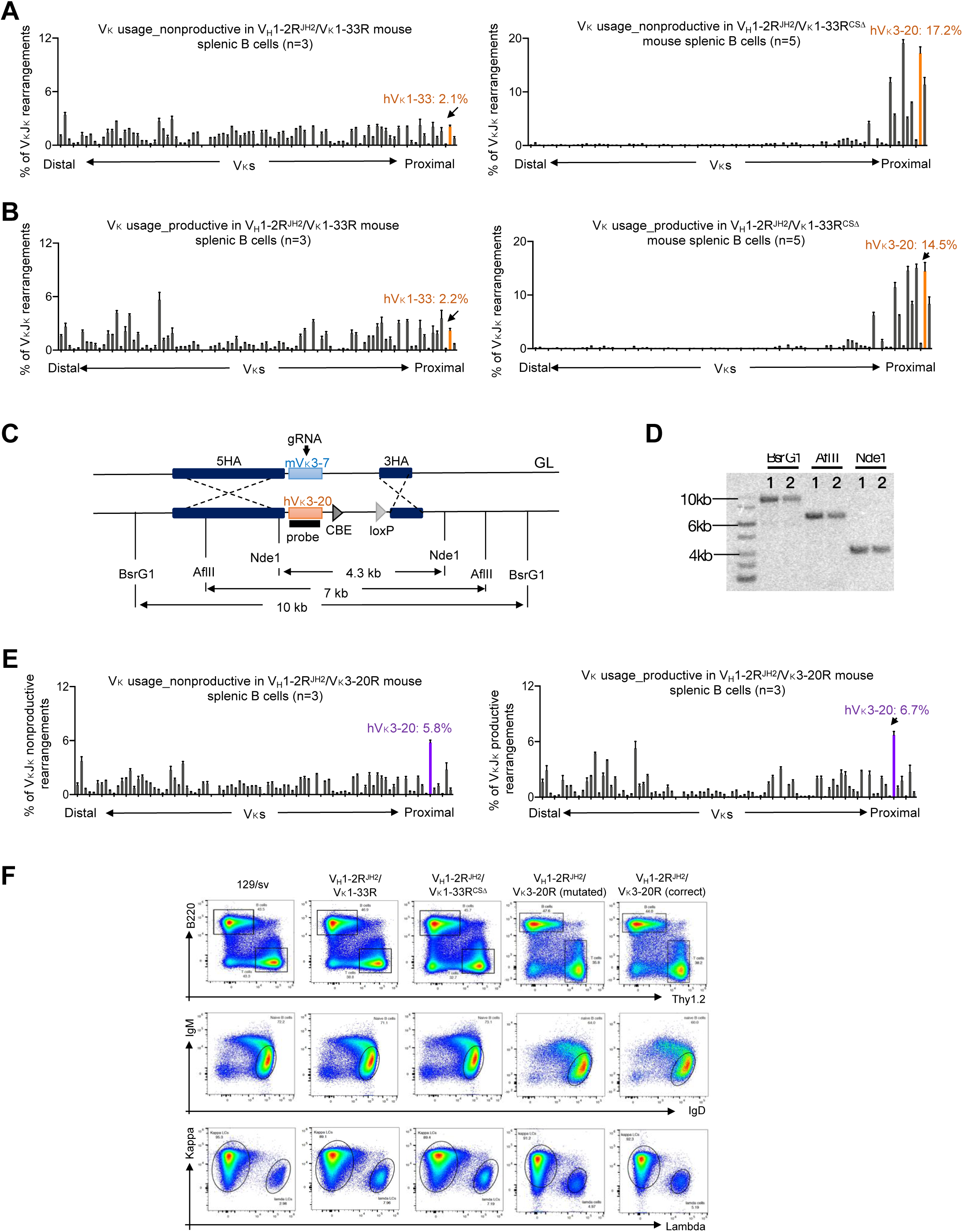
Generation and characterization of the human Vκ-rearranging light chains. (A) HTGTS-rep-seq analyses of Vκ nonproductive rearrangements in V_H_1-2R^JH2^/Vκ1-33R mouse splenic B cells (left) and V_H_1-2R^JH2^/Vκ1-33R^CSΔ^ mouse splenic B cells (right). The histogram displays the percent of nonproductive rearrangements of each Vκ among all nonproductive VκJκ rearrangements. The Vκ1-33 was labeled in orange. The percentage of Vκ segments in nonproductive rearrangements represents the V usage in primary V(D)J recombination. (B) HTGTS-rep-seq analyses of Vκ productive rearrangements in V_H_1-2R^JH2^/Vκ1-33R mouse splenic B cells (left) and V_H_1-2R^JH2^/Vκ1-33R^CSΔ^ mouse splenic B cells (right). The histogram displays the percent of productive rearrangements of each Vκ among all productive VκJκ rearrangements. The Vκ1-33 was labeled in orange. (C) The diagram, not drawn to scale, illustrates the restriction digests and Southern probe that were used to differentiate the region before (GL) and after Vκ3-20 replacement (Vκ3-20-rearranging allele). (D) Southern analysis of positive ES clones that showed in (C). (E) HTGTS-rep-seq analyses of Vκ nonproductive (left panel) or productive (right panel) rearrangements in V_H_1-2^JH2^/Vκ3-20-rearranging splenic B cells. The Vκ3-20 was labeled in purple. (F) FACS analyses of splenic B cells from wild-type 129/Sv, V_H_1-2R^JH2^/Vκ1-33R, V_H_1-2R^JH2^/Vκ1-33R^CSΔ^, V_H_1-2R^JH2^/Vκ3-20R (mutated) and V_H_1-2R^JH2^/Vκ3-20R (correct) mice. We repeated these analyses in 3 mice and they show similar results. Data from (A), (B) and (E) were average of >=3 experimental repeats with error bars representing SDs.

**Figure S4.**
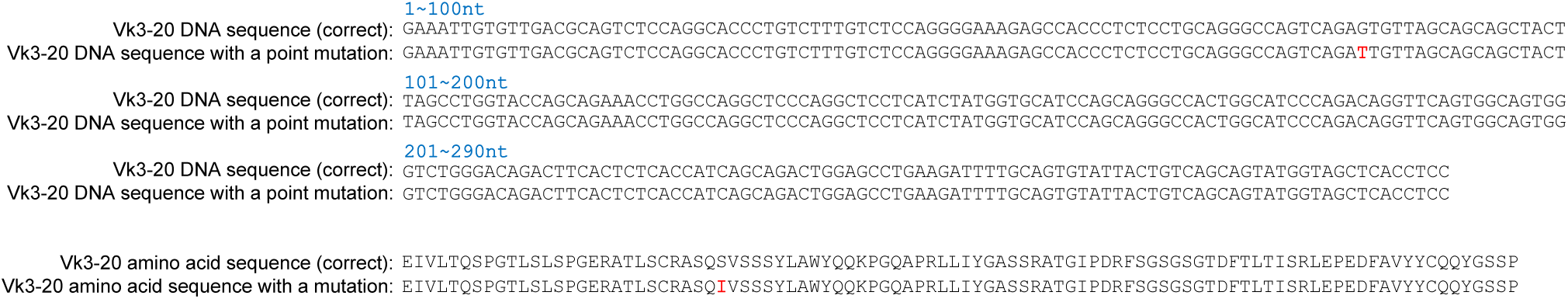
Mutation correction on the Vκ3-20 allele. A point mutation labeled in red on the nucleotides (upper) and amino acid (bottom) sequences of Vκ3-20 LC.

**Figure S5.**
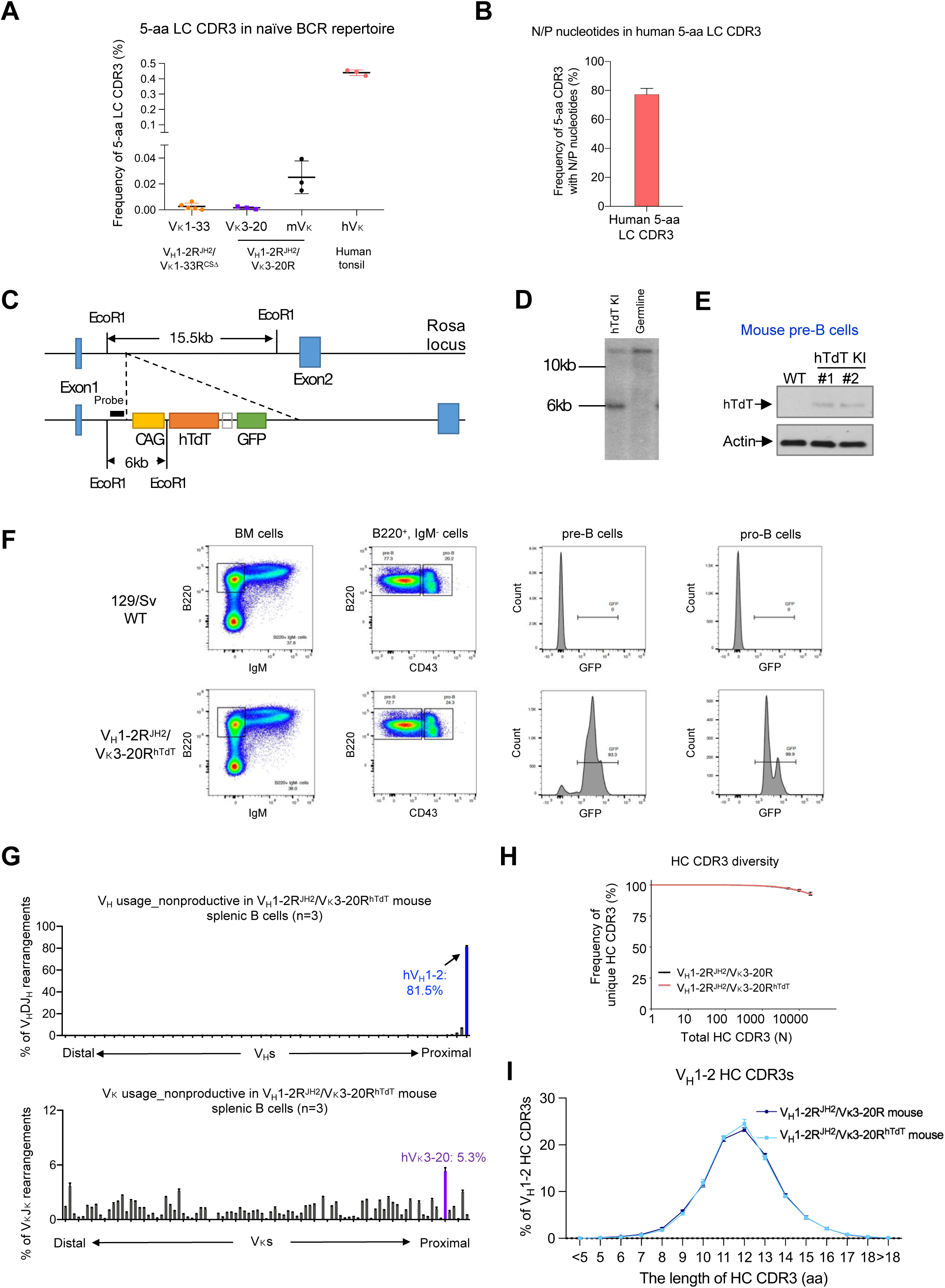
Enforced human TdT expression in the V_H_1-2^JH2^/Vκ3-20-rearranging mouse models. (A) Frequency of Vκ3-20, Vκ1-33, mouse Igκ and human Igκ LCs with 5-aa CDR3s in our VRC01-rearranging mouse splenic B cells and human tonsil naïve B cells. (B) Distribution of N or P nucleotides in human naïve Igκ LCs with 5-aa CDR3s. (C) The diagram illustrates the restriction digest and southern probe that were used to differentiate the region before and after human TdT knock-in. (D) Southern analysis of ES clone with hTdT knock-in. (E) Western Blot of TdT expression in mouse pre-B cells before and after hTdT knock-in. The TdT antibody can detect both human and mouse TdT. (F) FACS analyses of bone marrow B cells from 129/Sv wild-type and V_H_1-2R^JH2^/Vκ3-20R^hTdT^ mice. The pre-B cells were defined by B220^+^, IgM^-^ and CD43^-^. The pro-B cells were defined by B220^+^, IgM^-^ and CD43^+^. The GFP expression was linked with TdT expression as they shared a same promoter. (G) HTGTS-rep-seq analyses of nonproductive V_H_ (upper panel) or Vκ (bottom panel) usages in V_H_1-2R^JH2^/Vκ3-20R^hTdT^ mouse splenic B cells. The x axis represented V_H_ or Vκ locus from the distal to the *J*-proximal ends. The histogram displays the percent of usage of each V_H_ or Vκ among all nonproductive V_H_(D)J_H_ or VκJκ rearrangements. The usage of human V_H_1-2 was labeled in blue and the usage of human Vκ3-20 was labeled in purple. (H) The diversity of HC CDR3s in V_H_1-2R^JH2^/Vκ3-20R mouse and V_H_1-2R^JH2^/Vκ3-20R^hTdT^ mouse splenic B cells. The x axis represents the total HC CDR3 number (N). The y axis represents the frequency of unique HC CDR3s among total HC CDR3s. The differences of CDR3 diversities between V_H_1-2R^JH2^/Vκ3-20R mouse and V_H_1-2R^JH2^/Vκ3-20R^hTdT^ are not significant. (I) Length distribution of HC CDR3s in V_H_1-2R^JH2^/Vκ3-20R and V_H_1-2R^JH2^/Vκ3-20R^hTdT^ mouse splenic B cells. The differences measured by t-test were not significant. Data from (A), (B), (G) and (I) were mean ± SD of >=3 libraries from different mice.

**Figure S6.**
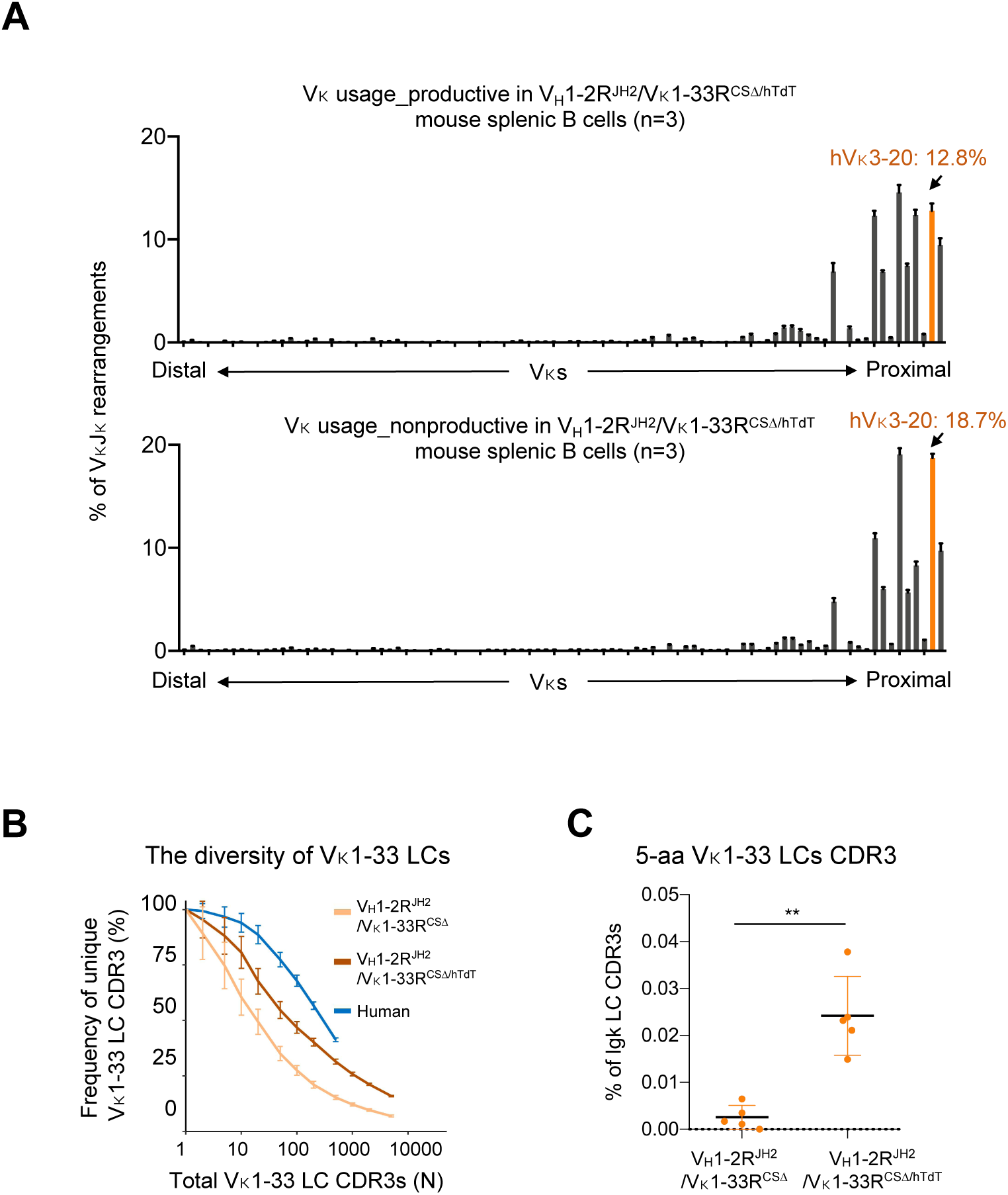
Enforced human TdT expression in the V_H_1-2^JH2^/Vκ1-33R^CSΔ^-rearranging mouse models. (A) HTGTS-rep-seq analysis of Vκ productive (upper panel) or nonproductive (bottom panel) rearrangements in V_H_1-2^JH2^/Vκ1-33^CSΔ/hTdT^-rearranging splenic B cells. The Vκ1-33 was labeled in orange. The percentage of Vκ segments in nonproductive rearrangements represents the V usage in primary V(D)J recombination. (B) The diversity of Vκ1-33 LC CDR3s in human, V_H_1-2R^JH2^/Vκ1-33R^CSΔ^ and V_H_1-2R^JH2^/Vκ1-33R^CSΔ/hTdT^ mouse naïve B cells. The differences of CDR3 diversities between V_H_1-2R^JH2^/Vκ1-33R^CSΔ^ and V_H_1-2R^JH2^/Vκ1-33R^CSΔ/hTdT^ mice are significant when the total CDR3 number is above 10 (*p*<0.001 for N>=10). (C) The frequency of 5-aa Vκ1-33 LC CDR3s in V_H_1-2R^JH2^/Vκ1-33R^CSΔ^ and V_H_1-2R^JH2^/Vκ1-33R^CSΔ/hTdT^ mouse naïve B cells.

**Figure S7.**
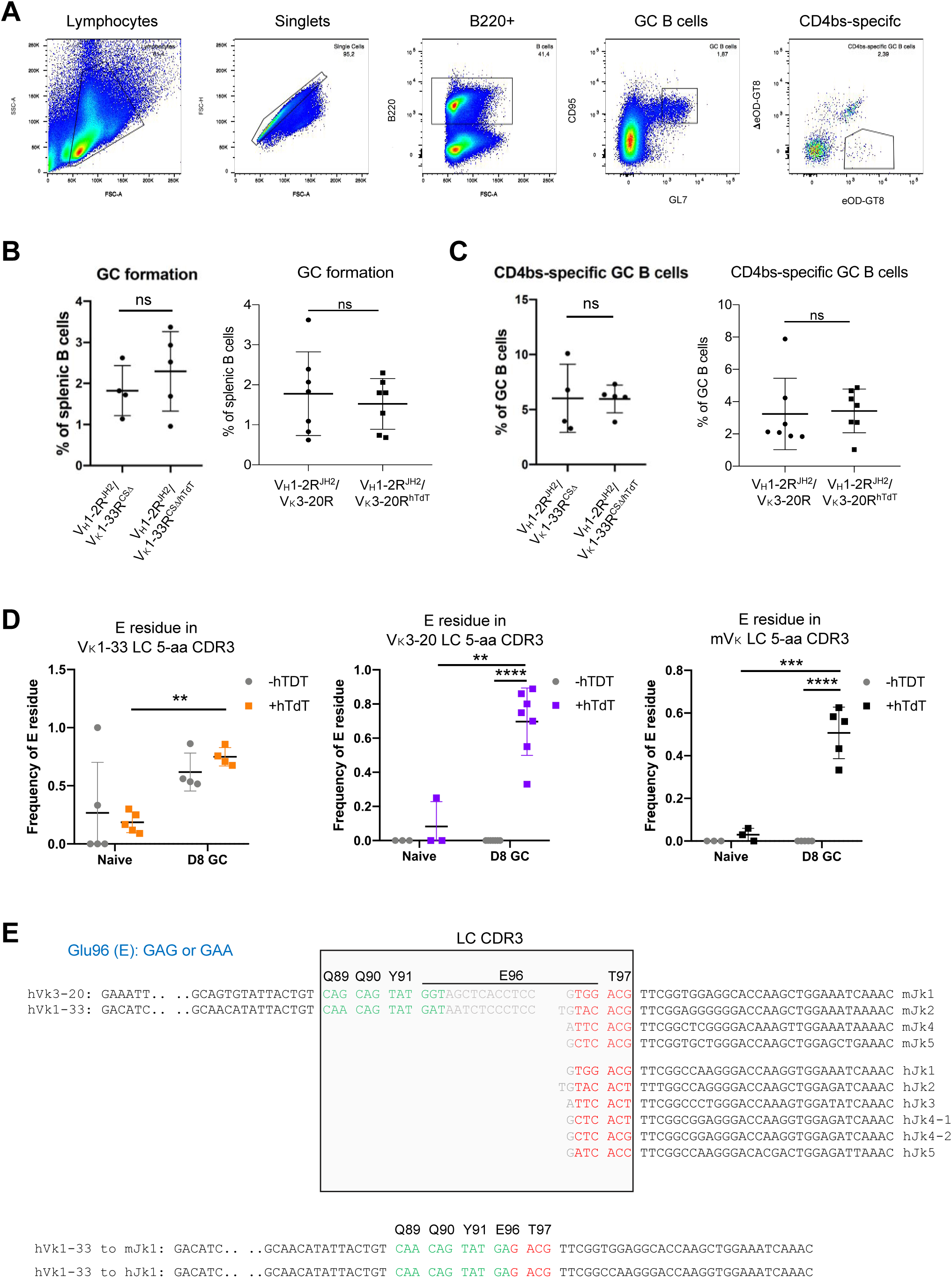
Human TdT enhanced VRC01-class GC responses induced by eOD-GT8 60mer, Related to Figure 2. (A) Gating strategy for single cell sorting of eOD-GT8-specific germinal center B cells after eOD-GT8 60mer immunization. (B) Proportion of GC B cells in V_H_1-2R^JH2^/Vκ1-33R^CSΔ^, V_H_1-2R^JH2^/Vκ3-20R, V_H_1-2R^JH2^/Vκ1-33R^CSΔ/hTdT^ and V_H_1-2R^JH2^/Vκ3-20R^hTdT^ mice. Each point represented one mouse. (C) Proportion of CD4bs-specific GC B cells in V_H_1-2R^JH2^/Vκ1-33R^CSΔ^, V_H_1-2R^JH2^/Vκ3-20R, V_H_1-2R^JH2^/Vκ1-33R^CSΔ/hTdT^ and V_H_1-2R^JH2^/Vκ3-20R^hTdT^ mice. Each dot represents one mouse. (D) The frequency of Glu96 (E) residue in 5-aa CDR3s of Vκ1-33, Vκ3-20 and mouse LCs before (naïve) and after eOD-GT8 60mer immunization (D8 GC). Each dot represents one mouse. (E) The Glu96 (E) residue formation in 5-aa CDRs of Vκ1-33, Vκ3-20 and mouse LCs. The Glu (E) amino acid is encoded by GAA or GAG. Both Vκ1-33 and Vκ3-20 can provide the G at the first position, but only Vκ1-33 can provide the A at the second position. On the other side, both mouse and human Jκs cannot provide the G and A at first and second positions, but mouse or human Jκ1 can provide G at the third position. Altogether, in the mouse pre-B cell lacking of TdT expression, the Glu96 (E) is formed when Vκ1-33 joins to mouse Jκ1. Other combinations failed to form the E residue. By examination of the mouse Vκ sequences, only Vκ14-111 can form the E residue in 5-aa LC CDR3 without N region added by TdT. But Vκ14-111 was not observed in the GCs induced by eOD-GT8, probably due to the low affinity of V region to eOD-GT8. Statistical comparisons in (**B**), (**C**) and (**D**) were performed using unpaired, two-tail t-test. \**p* <0.05, \*\**p* <0.01, \*\*\**p* <0.001, \*\*\*\**p* <0.0001

**Figure S8.**
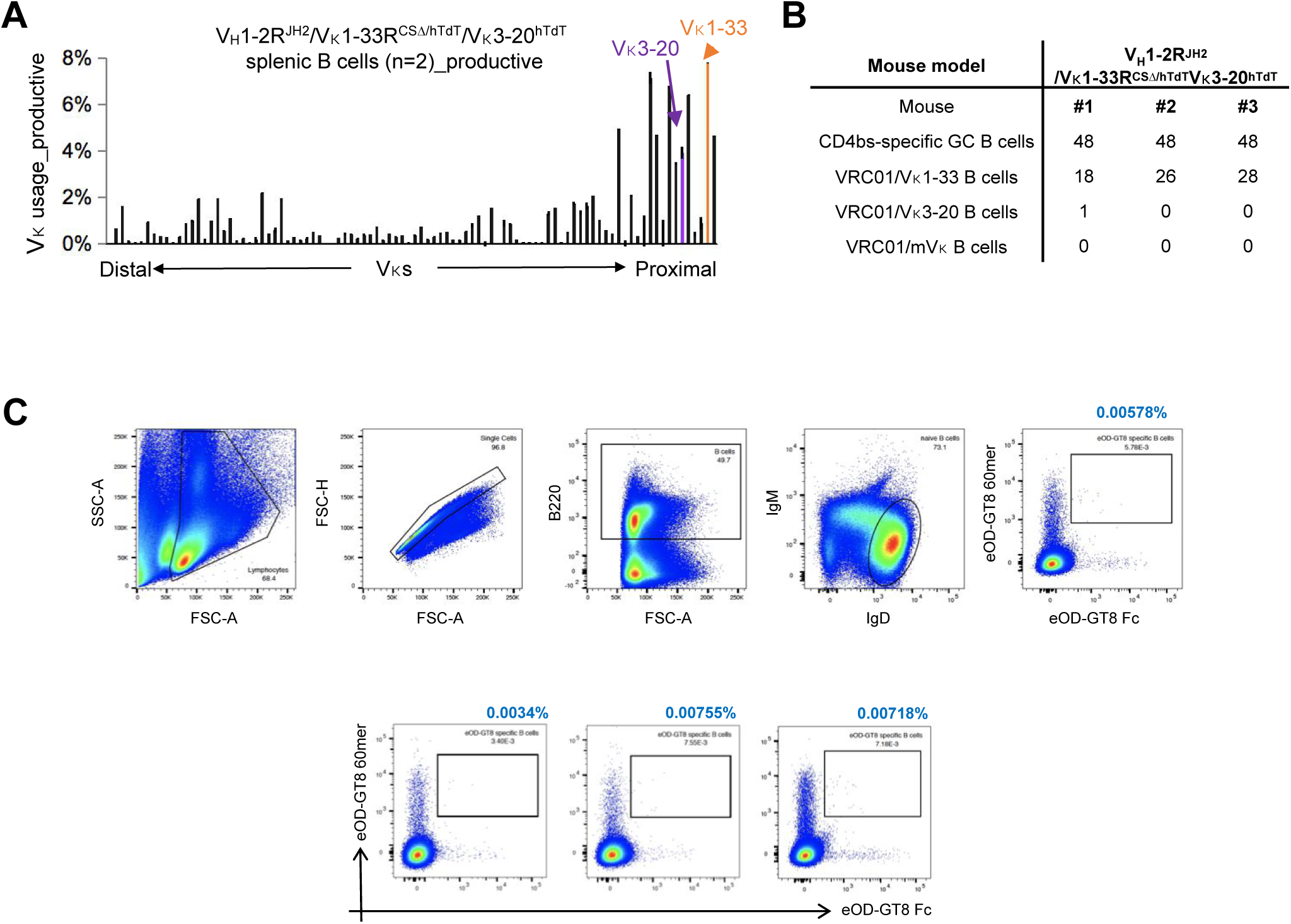
Generation and characterization of V_H_1-2^JH2^/Vκ1-33^CSΔ/hTdT^/Vκ3-20^hTdT^ and V_H_1-2^JH2^/Vκ1-33/Vκ3-20^hTdT^ -rearranging mouse models. (A) HTGTS-rep-seq analysis of Vκ usage in V_H_1-2R^JH2^/Vκ1-33R^CSΔ/hTdT^/Vκ3-20R^hTdT^ mouse splenic B cells. The usage of human Vκ1-33 is labeled in orange, and the usage of human Vκ3-20 is labeled in purple. (B) Table shown the VRC01-class B cells elicited by eOD-GT8 60mer in V_H_1-2R^JH2^/Vκ1-33R^CSΔ/hTdT^/Vκ3-20R^hTdT^ mice. 48 CD4bs-specific GC B cells were sorted from each mice on day 8 GCs post-immunization. The VRC01-class BCRs were identified by single cell RT-PCR following sanger sequencing. (C) Gating strategy for single cell sorting of eOD-GT8-specific naïve B cells in V_H_1-2^JH2^/Vκ1-33/Vκ3-20^hTdT^-rearranging mouse models.

**Figure S9.**
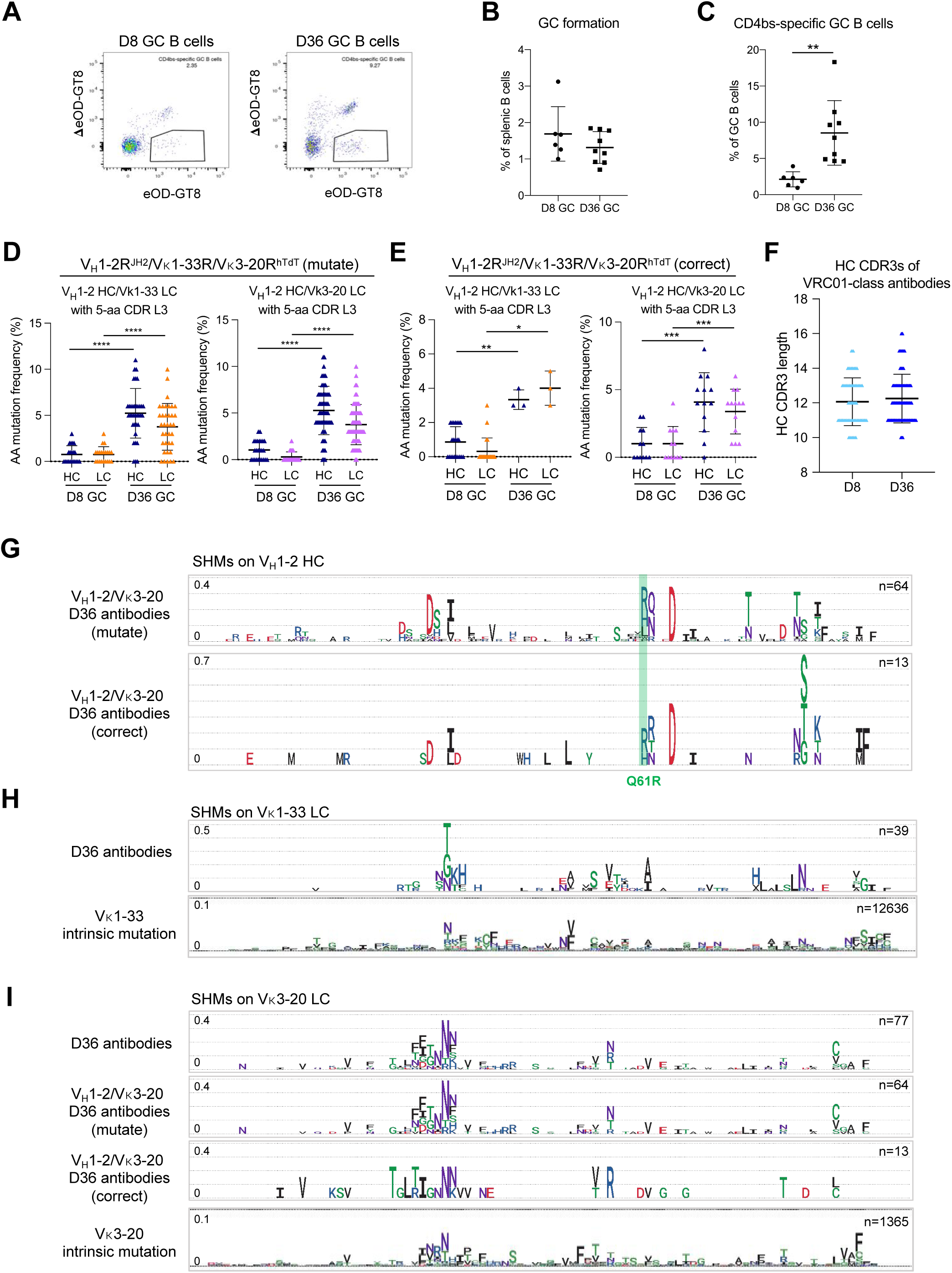
VRC01-class antibodies develop SHM and affinity maturation in GCs induced by eOD-GT8 60mer. (A) FACS analyses of GC B cells on both day 8 and day 36 post-immunization with eOD-GT8 60mer. The boxed CD4bs-specific GC B cells were sorted for single cell sequencing. (B) The Proportion of GC B cells in V_H_1-2R^JH2^/Vκ1-33R/Vκ3-20R^hTdT^ mice at day 8 and day 36 post-immunization. (C) Proportion of CD4bs-specific GC B cells in V_H_1-2R^JH2^/Vκ1-33R/Vκ3-20R^hTdT^ mice at day 8 and day 36 post-immunization. (D) Amino acid mutation in VRC01-class antibodies cloned from day 8 and day 36 GCs in V_H_1-2R^JH2^/Vκ1-33R/Vκ3-20R^hTdT^ mice with a germline mutation on Vκ3-20 allele. Each dot represents one HC or one LC. The median with interquartile range is plotted. (E) Amino acid mutation frequency in VRC01-class antibodies cloned from day 8 and day 36 GCs in V_H_1-2R^JH2^/Vκ1-33R/Vκ3-20R^hTdT^ mice with a correct Vκ3-20 allele. (F) Length distribution of HC CDR3s in all VRC01-class antibodies cloned from day 8 and day 36 GCs. (G) Mutation frequency of each amino acid on germline-encoded V_H_1-2 region of V_H_1-2/Vκ3-20 antibodies with (upper) or without (bottom) a germline mutation that cloned from day 36 GCs shown in sequence logo profiles. The distance between dotted horizontal lines representing 0.1 (10%). (H) Mutation frequency of each amino acid on germline-encoded Vκ1-33 region of VRC01-class antibodies that cloned from day 36 GCs shown in sequence logo profiles. The distance between dotted horizontal lines representing 0.1 (10%). For reference, the intrinsic mutation patterns from non-productive rearrangements are represented below. (I) Mutation frequency of each amino acid on germline-encoded Vκ3-20 region of V_H_1-2/Vκ3-20 antibodies that cloned from day 36 GCs shown in sequence logo profiles. 4 panels from top to bottom showed all V_H_1-2/Vκ3-20 antibodies, V_H_1-2/Vκ3-20 antibodies with a germline mutation, V_H_1-2/Vκ3-20 antibodies with correct sequences, and nonproductive Vκ3-20 sequences that represents the intrinsic mutation pattern. The distance between dotted horizontal lines representing 0.1 (10%). Statistical comparisons in (C), (D) and (E) were performed using a two-tailed unpaired t test. \**p* <0.05, \*\**p* <0.01, \*\*\**p* <0.001, \*\*\*\**p* <0.0001

**Table S1.**
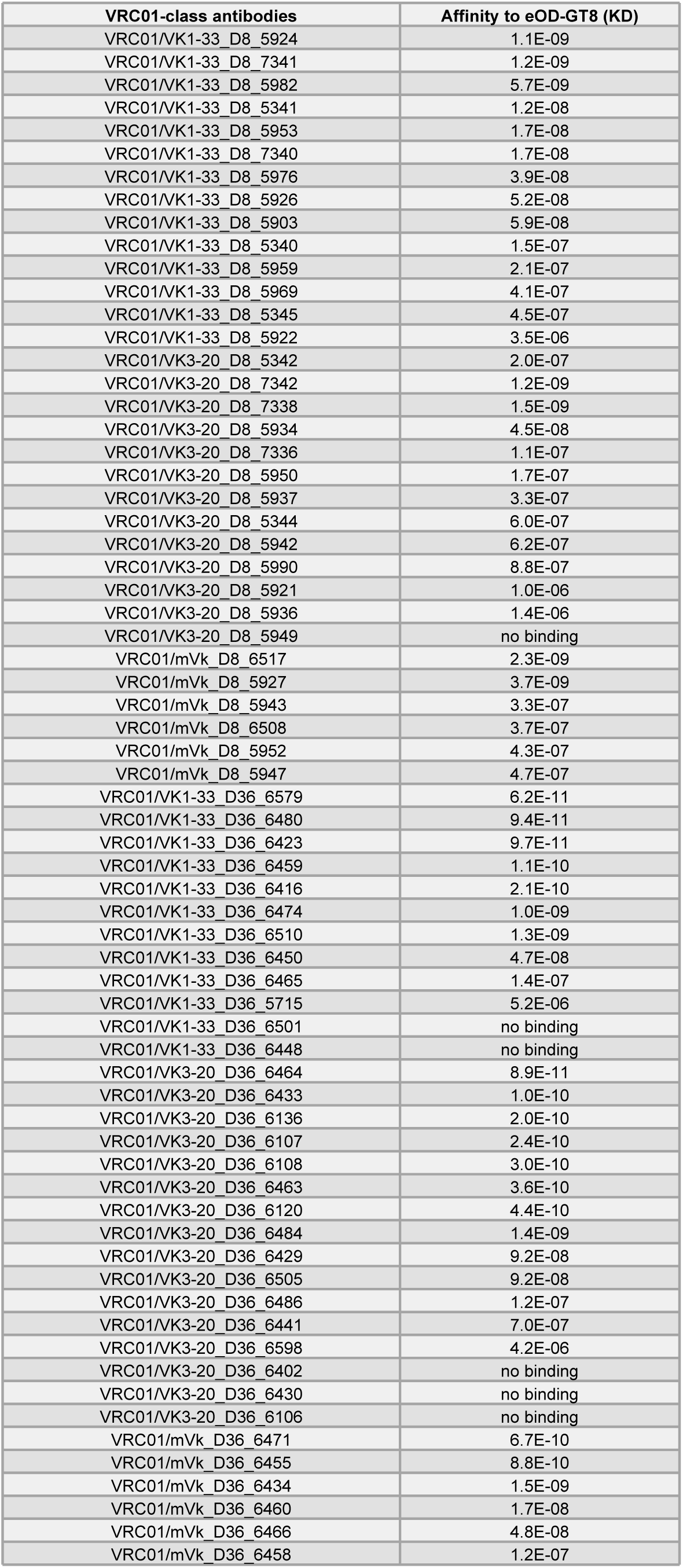
eOD-GT8 binding affinity of VRC01-class antibodies.

| **VRC01-class antibodies** | **Affinity to eOD-GT8 (KD)** |
| --- | --- |
| VRC01/VK1-33_D8_5924 | 1.1E-09 |
| VRC01/VK1-33_D8_7341 | 1.2E-09 |
| VRC01/VK1-33_D8_5982 | 5.7E-09 |
| VRC01/VK1-33_D8_5341 | 1.2E-08 |
| VRC01/VK1-33_D8_5953 | 1.7E-08 |
| VRC01/VK1-33_D8_7340 | 1.7E-08 |
| VRC01/VK1-33_D8_5976 | 3.9E-08 |
| VRC01/VK1-33_D8_5926 | 5.2E-08 |
| VRC01/VK1-33_D8_5903 | 5.9E-08 |
| VRC01/VK1-33_D8_5340 | 1.5E-07 |
| VRC01/VK1-33_D8_5959 | 2.1E-07 |
| VRC01/VK1-33_D8_5969 | 4.1E-07 |
| VRC01/VK1-33_D8_5345 | 4.5E-07 |
| VRC01/VK1-33_D8_5922 | 3.5E-06 |
| VRC01/VK3-20_D8_5342 | 2.0E-07 |
| VRC01/VK3-20_D8_7342 | 1.2E-09 |
| VRC01/VK3-20_D8_7338 | 1.5E-09 |
| VRC01/VK3-20_D8_5934 | 4.5E-08 |
| VRC01/VK3-20_D8_7336 | 1.1E-07 |
| VRC01/VK3-20_D8_5950 | 1.7E-07 |
| VRC01/VK3-20_D8_5937 | 3.3E-07 |
| VRC01/VK3-20_D8_5344 | 6.0E-07 |
| VRC01/VK3-20_D8_5942 | 6.2E-07 |
| VRC01/VK3-20_D8_5990 | 8.8E-07 |
| VRC01/VK3-20_D8_5921 | 1.0E-06 |
| VRC01/VK3-20_D8_5936 | 1.4E-06 |
| VRC01/VK3-20_D8_5949 | no binding |
| VRC01/mVk_D8_6517 | 2.3E-09 |
| VRC01/mVk_D8_5927 | 3.7E-09 |
| VRC01/mVk_D8_5943 | 3.3E-07 |
| VRC01/mVk_D8_6508 | 3.7E-07 |
| VRC01/mVk_D8_5952 | 4.3E-07 |
| VRC01/mVk_D8_5947 | 4.7E-07 |
| VRC01/VK1-33_D36_6579 | 6.2E-11 |
| VRC01/VK1-33_D36_6480 | 9.4E-11 |
| VRC01/VK1-33_D36_6423 | 9.7E-11 |
| VRC01/VK1-33_D36_6459 | 1.1E-10 |
| VRC01/VK1-33_D36_6416 | 2.1E-10 |
| VRC01/VK1-33_D36_6474 | 1.0E-09 |
| VRC01/VK1-33_D36_6510 | 1.3E-09 |
| VRC01/VK1-33_D36_6450 | 4.7E-08 |
| VRC01/VK1-33_D36_6465 | 1.4E-07 |
| VRC01/VK1-33_D36_5715 | 5.2E-06 |
| VRC01/VK1-33_D36_6501 | no binding |
| VRC01/VK1-33_D36_6448 | no binding |
| VRC01/VK3-20_D36_6464 | 8.9E-11 |
| VRC01/VK3-20_D36_6433 | 1.0E-10 |
| VRC01/VK3-20_D36_6136 | 2.0E-10 |
| VRC01/VK3-20_D36_6107 | 2.4E-10 |
| VRC01/VK3-20_D36_6108 | 3.0E-10 |
| VRC01/VK3-20_D36_6463 | 3.6E-10 |
| VRC01/VK3-20_D36_6120 | 4.4E-10 |
| VRC01/VK3-20_D36_6484 | 1.4E-09 |
| VRC01/VK3-20_D36_6429 | 9.2E-08 |
| VRC01/VK3-20_D36_6505 | 9.2E-08 |
| VRC01/VK3-20_D36_6486 | 1.2E-07 |
| VRC01/VK3-20_D36_6441 | 7.0E-07 |
| VRC01/VK3-20_D36_6598 | 4.2E-06 |
| VRC01/VK3-20_D36_6402 | no binding |
| VRC01/VK3-20_D36_6430 | no binding |
| VRC01/VK3-20_D36_6106 | no binding |
| VRC01/mVk_D36_6471 | 6.7E-10 |
| VRC01/mVk_D36_6455 | 8.8E-10 |
| VRC01/mVk_D36_6434 | 1.5E-09 |
| VRC01/mVk_D36_6460 | 1.7E-08 |
| VRC01/mVk_D36_6466 | 4.8E-08 |
| VRC01/mVk_D36_6458 | 1.2E-07 |

**Table S2.** VRC01-rearranging mouse model.

| Epitope | bnAb | Model | Human Ig segment | Other modification | hHC/LC % |
| --- | --- | --- | --- | --- | --- |
| CD4 binding site | VRC01 | $V_H1-2R^{JH2}/V_K1-33R^{hTdT}$ | hV<sub>H</sub>1-2, hJ<sub>H</sub>2, hV<sub>K</sub>1-33 | ΔIGCRI<br>hTdT | hHC(40%), hLC (2%) |
| | | $V_H1-2R^{JH2}/V_K1-33R^{CS\Delta/hTdT}$ | hV<sub>H</sub>1-2, hJ<sub>H</sub>2, hV<sub>K</sub>1-33 | ΔIGCRI, ΔCer/Sis,<br>hTdT | hHC(40%), hLC (13%) |
| | | $V_H1-2R^{JH2}/V_K3-20R^{hTdT}$ | hV<sub>H</sub>1-2, hJ<sub>H</sub>2, hV<sub>K</sub>3-20 | ΔIGCRI,<br>hTdT | hHC(40%), hLC (9%) |
| | | $V_H1-2R^{JH2}/V_K1-33R^{CS\Delta/hTdT}/V_K3-20R^{hTdT}$ | hV<sub>H</sub>1-2, hJ<sub>H</sub>2, hV<sub>K</sub>3-20,<br>hV<sub>K</sub>1-33 | ΔIGCRI,<br>ΔCer/Sis, hTdT | hHC(40%), hLC (hV<sub>K</sub>1-33:<br>7%; hV<sub>K</sub>3-20: 4%) |
| | | $V_H1-2R^{JH2}/V_K1-33R/V_K3-20R^{hTdT}$ | hV<sub>H</sub>1-2, hJ<sub>H</sub>2, hV<sub>K</sub>3-20,<br>hV<sub>K</sub>1-33 | ΔIGCRI,<br>hTdT | hHC(40%), hLC (hV<sub>K</sub>1-33:<br>1%; hV<sub>K</sub>3-20: 5%) |

**Table S3.**
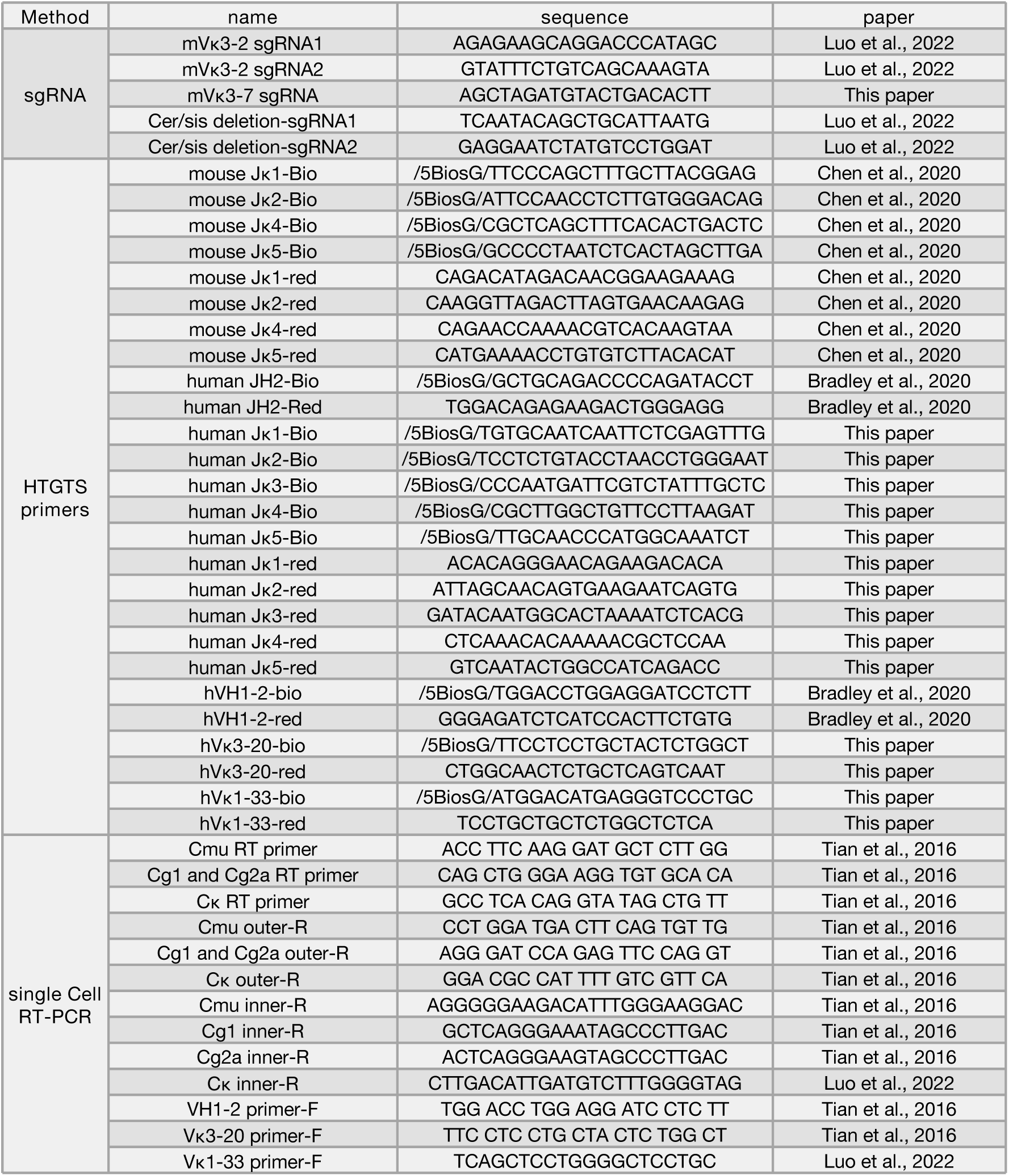
Primer sequences.

